# *Id1* Promotes Clonal Hematopoiesis in Mice with *Tet2* Loss of Function

**DOI:** 10.1101/2024.11.19.624318

**Authors:** Shweta Singh, Tanmoy Sarkar, Kristbjorn O. Gudmundsson, Brad L. Jakubison, Holly M. Morris, Sandra Burkett, Karim Baktiar, Gary T. Pauly, Dina M. Sigano, Joel P. Schneider, Jonathan R. Keller

## Abstract

Hematopoietic malignancies emerge through the gradual acquisition of genetic mutations within hematopoietic stem and progenitor cells (HSPCs). Mutations that occur early in disease progression impart a selective growth advantage to HSPCs, which allows them to expand and contribute to a substantial percentage of mature blood cells. This increased expansion is termed clonal hematopoiesis (CH) and is a preleukemic phase associated with an increased risk of developing leukemia. Inhibitor of DNA binding 1 (ID1) protein is a transcriptional regulator of proliferation/differentiation of neuronal, muscle, hematopoietic and other cells, and is frequently overexpressed in cancer. *Id1* is expressed at low levels in normal HSCs and is induced by growth factors and other mediators of inflammatory stress and promotes HSPC proliferation *in vitro and in vivo.* Since chronic inflammation is associated with the progression of hematopoietic malignancies, reducing *Id1* expression during CH may be therapeutic. Mutations in *TET2* are frequently observed in patients with CH, and *Tet2^−/−^*mice develop CH that progress to hematopoietic malignancies. *Id1* is upregulated in murine *Tet2^−/−^* HSPCs and in AML, CMML and MDS patient samples with *TET2* mutations. Genetic ablation of *Id1* in *Tet2^−/−^* HSPCs reduces HSPC expansion/self-renewal/CH, extramedullary hematopoiesis, myeloid skewing, genetic instability and delays the onset of disease. Mechanistically, p16 expression, senescence and apoptosis were increased and proliferation decreased in *Tet2^−/−^; Id1^−/−^* HSPCs. Thus, ID1 may represent a potential therapeutic target to reduce CH, hematopoietic hyperplasia, and delay the onset of disease.

**One Sentence Summary:** Genetic ablation of *Id1* in *Tet2^−/−^* mice rescues clonal hematopoiesis by increasing CDKI expression, apoptosis, senescence, and differentiation, and reducing cell growth.

## INTRODUCTION

Hematopoietic malignancies arise, in part, from the accumulation of somatic mutations in HSPCs as individual age (*1–3*). Mutations that occur early in disease progression can impart an increased fitness/survival advantage to HSPCs, which allows them to expand relative to normal HSPCs. Consequently, during the early preleukemic stages of hematopoietic malignancies the frequency of mature blood cells with these founding mutations is increased in patients without clinical signs of hematologic dysfunction, a condition known as clonal hematopoiesis (CH)(*4–6*). Patients with CH have a roughly ten-fold increased frequency of progressing to hematopoietic malignancies via accumulation of additional mutations during aging, indicating the need for therapeutic interventions during this early preleukemic phase of the malignancy(*1, 4, 7*). Loss of function mutations in the epigenetic modifying enzyme, ten-eleven translocation 2 (*TET2),* are frequently observed in patients with CH, myeloproliferative neoplasms (MPN), chronic myelomonocytic leukemia (CMML), systemic mastocytosis, myelodysplastic syndrome (MDS) and acute myeloid leukemia (AML)(*8–13*). Mice with *Tet2* loss of function (*Tet2^−/−^*) show CH that progresses to myeloid and lymphoid hyperplasia, and leukemia in some mice after a long latency(*14–22*). *Tet2^−/−^* mice are characterized by hyperinflammatory environments with increased production of proinflammatory molecules including IL-6, IL-1, TNF-α and others, which contribute to HSPC expansion and artherosclerosis(*21, 23–28*). Inhibition of inflammatory cytokines, their receptors and downstream signaling proteins reduces HSPC expansion and CH in *Tet2^−/−^* mice suggesting that cytokines drive preleukemic HSPC expansion(*23, 27, 29, 30*). Chronic inflammation has been implicated in promoting the progression of myeloid malignancies and suggest suggesting that reducing inflammation could slow the progression of hematopoietic malignancies(*31–35*).

Inhibitor of DNA binding proteins (ID1-4) are helix-loop-helix transcription factors that regulate the proliferation and differentiation of neuronal, muscle, endothelial, hematopoietic and cells in other tissues, and are frequently deregulated in cancer(*36–38*). The founding member of this family, ID1, is expressed at low levels in HSCs and is induced by hematopoietic growth factors (HGFs) that promote myeloid proliferation and differentiation including IL-3 and GM-CSF(*39, 40*). Enforced expression of *Id1* in HSPCs enhances HGF-induced proliferation and immortalizes HPCs *in vitro* and promotes a myeloproliferative disease after a long latency *in vivo*(*41, 42*). *Id1* is induced in HSPC during BMT, genotoxic and inflammatory stress and aging *in vivo*(*43*). Since *Id1* is induced in HSPCs by proinflammatory cytokines *in vitro* and *in vivo,* and during BMT and inflammatory stress, and promotes HSPC proliferation, we reasoned that ID1 may represent a potential therapeutic target to reduce CH(*39, 41, 44*). We show here that ablation of *Id1* in *Tet2^−/−^* mice 1) reduces expansion/self-renewal of HSPCs, 2) reduces extramedullary hematopoiesis, 3) reduces myeloid skewing, 4) reduces genomic instability and 5) delays the onset of hematopoietic malignancy. Thus, inhibiting ID1 in patients with CH and chronic inflammation could be therapeutic and reduce HSPC expansion, MPN and the onset of hematopoietic disease.

## RESULTS

### *Id1* expression is increased in *TET2^−/−^* HSPCs and intrinsically promotes the expansion of hematopoietic stem and progenitor cells

Analysis of previously published RNA-seq data sets demonstrate that *Id1* RNA is increased in *Tet2^−/−^* HSPCs compared to *Tet2^+/+^* controls (Figure S1A)(*20, 45*). To confirm that *Id1* expression is increased in *Tet2^−/−^*HSPCs *in vivo*, we crossed *Tet2^−/−^* mice with a *Id1-GFP* reporter mouse model and found that ID1-GFP expression is significantly elevated in HSCs, MPPs and HPCs in *Tet2^−/−^;Id1^GFP/+^* mice (Figure 1A)(*46*). Furthermore, *Id1* mRNA is upregulated in FACS purified donor *Tet2^−/−^* HSPCs from competitively transplanted recipient mice (Figure 1B). While gene expression data sets of HSPCs from patients with CH and *TET2* mutations are currently limiting, *ID1* expression is increased in AML patient samples with TET2 mutations, and in CD34+ HSPCs from patients with MDS and CMML and *TET2* mutations compared to controls (Figure S1B-D). Collectively, these data demonstrate that *Id1* expression is increased in murine *Tet2^−/−^*HSPCs and in AML, MDS and CMML patient samples with *TET2* mutations.

**Fig. 1.**
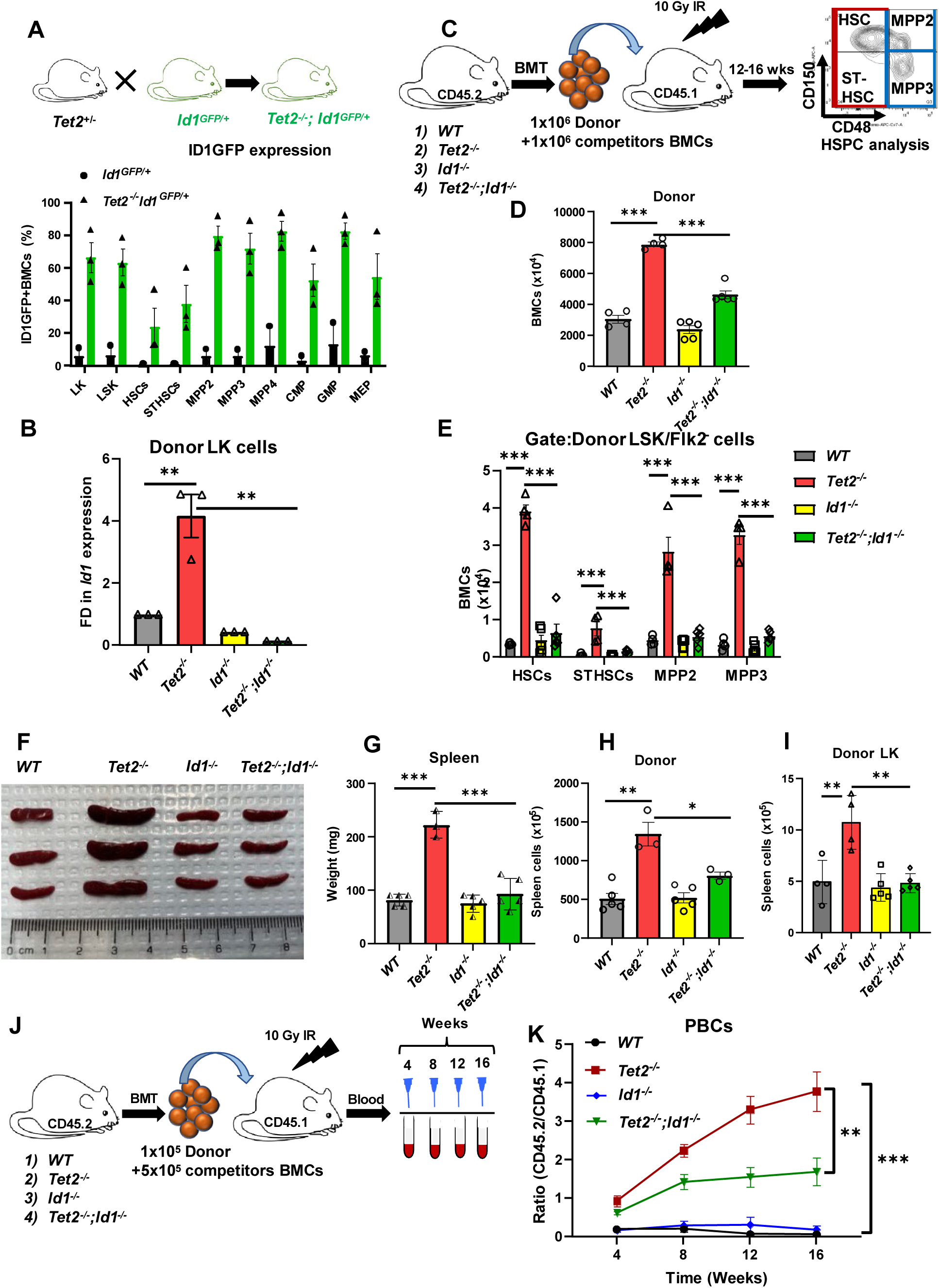
*Id1* ablation reduces the expansion of *Tet2^−/−^* HSPCs in BMT recipient mice. (A) Percentage ID1-GFP expression in HSPCs isolated from *Tet2^−/−^;Id1^GFP/+^ Tet2^+/+^;Id1^GFP/+^* mice by flow cytometry. (B) *Id1* expression in FACS purified donor derived LK cells isolated from competitive BMT recipients after 12 wks. (Fold Difference, FD). (C) Summary of experimental design for competitive bone marrow transplantation. (D) Total number of donor derived BMCs in competitive BMT recipient mice. (E) Total number of donor derived HSPCs in BM of primary recipient mice. (F) Photograph of spleens and (G) spleen weight in competitive BMT recipient mice. (H) Total number of donor derived and (I) donor derived LK cells in the spleens of competitive BMT recipient mice. (J) Summary of experimental design to detect clonal expansion. (K) Ratio of CD45.2/CD45.1 expression in PBCs of recipient mice 4-, 8-, 12– and 16-weeks after BMT. Data are presented as the mean ± SEM, and comparisons between mean values of groups were evaluated using an unpaired, 1-tailed Student’s t test. *\*P* ≤ 0.05, *\*\*P* ≤ 0.01 and *** *P* ≤ 0.001.

To investigate if *Id1* intrinsically contributes to CH we evaluated hematopoietic development in competitive repopulation assays to rule out any contribution that loss of *Tet2* function might have in the hematopoietic microenvironment (HME). Furthermore, γ-irradiation (γ-IR) conditioning for BMT induces pro-inflammation that promotes murine HSPC expansion, and CH is increased in human donor HSPCs after allogeneic BMT (*47–49*). *Tet2^−/−^*BMCs have a competitive repopulation advantage *in vivo*(*15, 17, 19*); therefore, we transplanted equal numbers of *Tet2^+/+^;Id1^+/+^(WT), Tet2^+/+^; Id1^−/−^ (Id1^−/−^), Tet2^−/−^*; *Id1^+/+^ (Tet2^−/−^)* and *Tet2^−/−^; Id1^−/−^* donor BMCs with host BMCs to evaluate if reducing *Id1* levels in *Tet2^−/−^*BMCs could affect donor reconstitution (Figure 1C). As expected, we observed an increase in the total number of donor-derived (CD-45.2+) *Tet2^−/−^* BMCs in primary recipient mice 12-16 weeks after BMT compared to mice transplanted with *Tet2^+/+^ and Id1^−/−^* BMCs (Figure 1D). *Tet2^−/−^; Id1^−/−^* BMCs showed reduced donor BM repopulation indicating that *Id1* is intrinsically required for the enhanced repopulation and expansion of *Tet2^−/−^* BMCs. Immunophenotype analysis of repopulating cells revealed increased numbers of donor-derived lineage-negative, Sca-1-postive, c-Kit-positive (LSK) cells, LSK/Flk2-neg (Flk2-) and multipotent progenitor-4 (MPP-4) (Figure S1E and Figure S1G), and more primitive HSCs, ST-HSCs, and MPPs in repopulating *Tet2^−/−^* BMCs compared to controls (Figure 1E). These HSPC progenitors were significantly reduced and rescued in mice transplanted with *Tet2^−/−^; Id1^−/−^* donor BMCs demonstrating that *Id1* is required for the expansion of immunophenotypic *Tet2^−/−^* HSPCs.

*Tet2^−/−^* mice show extramedullary hematopoiesis(*15, 17, 19*); therefore, we analyzed donor hematopoietic repopulation in spleens and confirmed that mice competitively transplanted with *Tet2^−/−^* BMCs show increased size, weight and total spleen cell numbers compared to controls (Figure 1F-G and Figure S1F), which were reduced and rescued in mice transplanted with *Tet2^−/−^; Id1^−/−^* BMCs. Increased numbers of donor and donor-derived LK cells (Figure 1H-I and Figure S2A) were observed in the spleens of mice transplanted with *Tet2^−/−^*, which were reduced and rescued in mice transplanted with *Tet2^−/−^; Id1^−/−^* donor BMCs demonstrating that *Id1* intrinsically promotes extramedullary hematopoiesis in *Tet2^−/−^* chimeric mice.

To confirm that *Id1* contributes to CH in *Tet2^−/−^* mice, we employed an experimental model of CH(*23*); where host cells were competitively transplanted in 5-fold excess to donor-derived experimental cells into γ-IR recipient mice (Figure 1J). The frequency of donor *Tet2^−/−^* BMC reconstitution (Donor/Host ratio) was increased 3-4-fold in PBCs over time, while the frequency of *Tet2^+/+^* and *Id1^−/−^* BMC reconstitution was less than 0.5-fold over the 16 weeks of analysis (Figure 1K). In comparison, transplantation of *Tet2^−/−^* HSPCs that lack *Id1* expression showed significantly reduced repopulation potential (1.5-fold) compared to *Tet2^−/−^* BMCs indicating that *Id1* contributes to CH of pre-leukemic *Tet2^−/−^* HSPCs.

### Ablation of *Id1* in *Tet2^−/−^* HSPCs reduces myeloid skewing *in vitro* and *in vivo*

To investigate if *Id1* intrinsically regulates myeloid skewing we evaluated lineage development in mice noncompetitively transplanted mice with *Tet2^+/+^, Id1^−/−^, Tet2^−/−^* and *Tet2^−/−^; Id1^−/−^*BMCs (Figure S2B). CBC analysis of PBCs 12-16 weeks after transplantation of *Tet2^−/−^*BMCs showed elevated levels of white blood cells, neutrophils, and monocytes (Figure S2C), which were reduced and rescued in mice transplanted with *Tet2^−/−^; Id1^−/−^* BMCs. Similarly, donor-derived BM neutrophils, monocytes, macrophages, and dendritic cells were increased in mice transplanted with *Tet2^−/−^* BMCs (Figure S2D and Figure S3A), while donor CD19^+^ B cells and Ter-119+ erythroid cells were decreased (Figure S3B). Mice transplanted with *Tet2^−/−^; Id1^−/−^* BMCs showed a decrease and rescue of all myeloid lineage cells in the BM (Figure S3A), while CD19^+^ B cells and erythroid cells were increased (Figure S3B). Mice transplanted with *Tet2^−/−^* BMCs showed increased donor myeloid cell repopulation in the spleen, and myeloid skewing in the spleen was reduced and rescued in mice transplanted with *Tet2^−/−^; Id1^−/−^* BMCs (Figure S3C-D). Finally, *Tet2^−/−^* Lin-BMCs showed increased numbers of differentiating monocytes and macrophages after six days culture compared to control Lin-BMCs, but was reduced and rescued in *Tet2^−/−^; Id1^−/−^* Lin-cells (Figure S3E). Collectively, these results suggest that *Id1* intrinsically promotes myeloid development and skewing in *Tet2^−/−^* mice. In agreement with this conclusion, forced expression of *Id1* in normal HSPCs promotes myeloid and inhibits lymphoid development *in vitro* and *in vivo*(*39, 40*).

We show here that donor-derived Lin-/c-Kit^+^ (LK) and committed progenitors (CMP and GMP) are increased in the BM (Figure 2A-B) and spleen (Figure S3F-G) of *Tet2^−/−^* chimeric mice. LK, CMP and GMP cells were reduced and rescued in the BM (Figure 2A-B) and spleen (Figure S3F-G) of *Tet2^−/−^; Id1^−/−^*chimeric mice, suggesting that *Id1* contributes to myeloid skewing in *Tet2^−/−^* mice by promoting myeloid progenitor cell expansion.

**Fig. 2.**
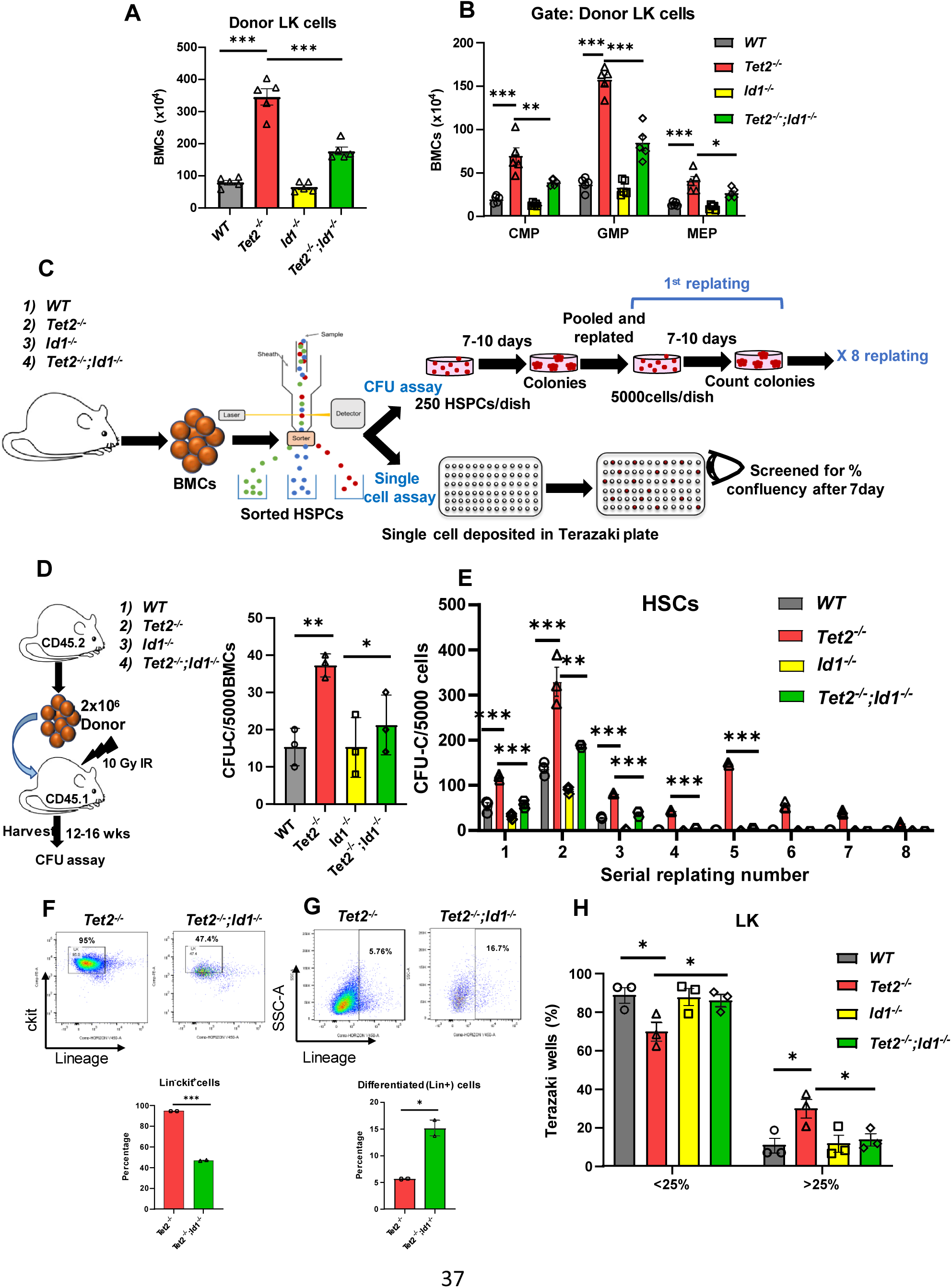
*Id1* promotes the expansion of *Tet2^−/−^* HSPCs *in vitro.* (A) Total number of donor derived LK and (B) CMP, GMP, and MEP cells in competitive BMT recipient mice. C) Summary of experimental design of self-renewal colony formation assay (CFU-c) and growth in single cell assays of FACS sorted HSPCs. (D) Number of CFU-c progenitors in 5 x 10^3^ BMCs from noncompetitive BMT recipient mice. (E) Number of CFU-c progenitors in 250 sorted HSCs in primary colony formation assays and, the number of CFU-c in 5 x 10^3^ serially replated cells in secondary and subsequent CFU-c assays. (F) Flow cytometry contour plots and percentages of LK cells and Lin^+^ cells present in pooled colonies from the 4^th^ serial replating assay of HSCs. (H) Single cell growth assay of FACS sorted LK cells in Terasaki plates representing percentage of wells with cell growth covering <25% or >25% of the well after seven days of culture. Data are presented as the mean ± SEM, and comparisons between mean values of groups were evaluated using an unpaired, 1-tailed Student’s t test. *\*P* ≤ 0.05, *\*\*P* ≤ 0.01 and *** *P* ≤ 0.001.

### *Id1* promotes progenitor expansion and self-renewal of *Tet2^−/−^*HSPCs by inhibiting cell differentiation

To confirm that *Id1* is critical for the expansion of functional *Tet2^−/−^* HSPCs, we examined HSPC function in colony formation (CFU-c), self-renewal and single cell growth assays (Figure 2C). BMCs from *Tet2^−/−^* chimeric mice showed increased colony formation compared to control mice, and the number of colonies were decreased and rescued in *Tet2^−/−^; Id1^−/−^* BMCs (Figure 2D). Thus, *Id1* is required for the expansion of functional HSPCs in *Tet2^−/−^*chimeric mice *in vivo*.

*Tet2^−/−^* HSPCs have enhanced self-renewal potential *in vitro*(*17*); therefore, we evaluated the replating potential of 5 x 10^3^ cells from CFU-c plates initiated from 250 FACS sorted HSCs (Figure 2E). As predicted, *Tet2^−/−^* BMCs showed extensive replating potential in CFU-c assays, while control BMCs showed limited replating potential that was significantly reduced after the third and fourth replating. *Tet2^−/−^*BMCs that lack *Id1* lost self-renewal potential indicating that *Id1* is required for the self-renewal of *Tet2^−/−^* BMCs (Figure 2E). We found that 90-95% of the cells in *Tet2^−/−^* colonies harvested from the 4^th^ serial replating were (Lin-/c-kit+, LK-like) while 40-45% of the cells in *Tet2^−/−^*; *Id1^−/−^* colonies were LK-like (Figure 2F). Further analysis revealed that cells in *Tet2^−/−^*; *Id1^−/−^* colonies showed increased numbers of differentiated Lin+ myeloid cells compared to cells *Tet2^−/−^*colonies suggesting that *Tet2^−/−^* HSPCs had undergone increased differentiation in the absence of *Id1* (Figure 2G). Thus, ID1 may contribute to the self-renewal of *Tet2^−/−^* cells by preventing their differentiation. In agreement with these observations, enforced expression of *Id1* in normal HSPCs promotes proliferation, prevents differentiation, and immortalizes SCF-dependent LK-like HSPCs *in vitro,* that further differentiate in response to HGFs in vitro and *in vivo*(*41*).

The requirement for *Id1* in the expansion of *Tet2^−/−^*HSPCs was confirmed by direct examination of single cell growth in Terasaki plates (20 ul/well) by bright field examination. Single *Tet2^−/−^* LK cells showed a significant reduction in wells with cell growth covering <25% of the well, and an increase in wells with cell growth covering >25% of the well compared to controls indicating that *Tet2^−/−^* LK cells have increased growth potential (Figure 2H). Ablation of *Id1* in *Tet2^−/−^* LK cells reduced and rescued cell growth resulting in a decrease in wells with cell growth >25% and increase in wells with cell growth <25% of the well. Similar results were observed in single cell assays with *Tet2^−/−^* HSCs that have a higher growth potential overall in this assay (Figure S2H). Thus, *Id1* promotes the expansion of *Tet2^−/−^* HSPCs *in vitro*.

### *Id1* inhibits apoptosis and senescence of *Tet2^−/−^* HSPCs

To understand how *Id1* contributes to HSPCs expansion we examined cell cycling, proliferation, apoptosis, and senescence of HSPCs in *Tet2^−/−^* chimeric mice. *Tet2^−/−^* MPP BMCs in chimeric mice showed increased BrdU incorporation compared to control cells that was reduced and rescued in *Tet2^−/−^*; *Id1^−/−^*MPPs, while *Tet2^−/−^* HSCs/STHSCs from the same mice did not show increased BrdU incorporation compared to controls (Figure S4A-B). In comparison, splenic *Tet2^−/−^* HSCs/STHSCs/MPP2s showed significantly increased in BrdU incorporation compared to controls, and loss of *Id1* expression reduced and rescued the proliferation of spleen HSPCs (Figure S4C). Overall, ID1 promotes the proliferation of *Tet2^−/−^* spleen and to a lesser extent BM HSPCs.

Analysis of apoptosis in *Tet2^−/−^* BMCs mice showed a decrease in Annexin+ apoptotic and increased numbers of live HSCs, ST-HSCs and MPPs compared to controls (Figure 3A-B). Loss of *Id1* expression in *Tet2^−/−^* HSPCs resulted in increased apoptosis and reduced the number of live cells. These results were confirmed in Lin-expansion assays *in vitro,* where *Tet2^−/−^* MPPs showed reduced apoptosis and increased viability compared to controls, while *Tet2^−/−^*; *Id1^−/−^*MPPs showed increased apoptosis and decreased survival (Figure 3C-E). Tunnel assays confirmed that *Tet2^−/−^* LK cells from chimeric mice show reduced apoptosis compared to control cells, while apoptosis was rescued and increased in *Tet2^−/−^*; *Id1^−/−^* LK cells (Figure 3F-H). Reduced levels of apoptosis have been observed in *Tet2^−/−^* Kit^+^ progenitors, and in *Tet2^−/−^* HSPCs in studies examining the mechanistic role of IL-6 in CH(*16, 23*). Taken together, these results suggest that *Id1* contributes to the expansion of *Tet2^−/−^* BM HSPCs by inhibiting apoptosis and promoting cell survival.

**Fig. 3.**
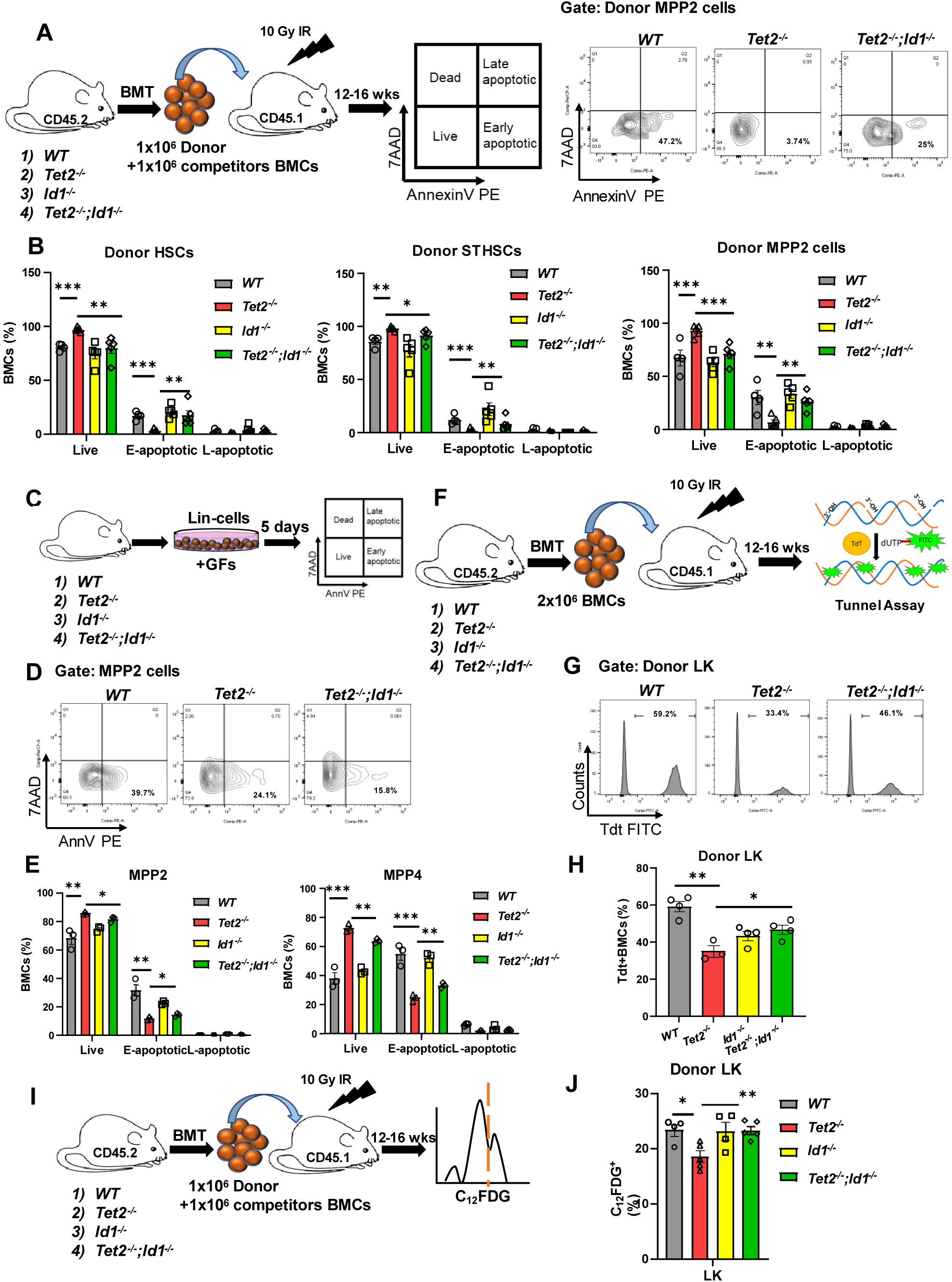
*Id1* inhibits apoptosis and senescence of *Tet2^−/−^* HSPCs. (A) Summary of **e**xperimental design to evaluate apoptosis by Annexin V staining in donor derived HSPCs from competitive BMT recipient mice and flow cytometry contour plots of Annexin V and 7AAD stained donor derived MPP2 BMCs from the indicated recipient mice. (B) Percentage of live, early (E) and late (L) apoptotic cells in donor derived HSCs, STHSCs and MPPs from chimeric mice by flow cytometry. (C) Summary of experimental design to evaluate apoptosis of Lin^−^ BMCs cultured *in vitro*. (D) Flow cytometry contour plots of Annexin V and 7AAD stained MPP2 cells from *in vitro* cultures after 5 days. (E) Percentage of live, E– and L-apoptotic MPP2 and MPP4 cells from *in vitro* cultures. (F) Summary of experimental design to evaluate apoptosis by tunnel assay in HSPCs from noncompetitive BMT recipient mice. (G) Flow cytometry histograms of Tdt^+^ donor derived LK BMCs from noncompetitive BMT recipient mice. (H) Percentage of donor derived Tdt^+^ donor derived LK cells in BMT recipient mice. (I) Summary of experimental design to detect senescent cells in competitive BMT recipient mice by flow cytometry. (J) Frequency of C_12_FDG^+^ donor derived LK cells in chimeric mice. Data are presented as the mean ± SEM, and comparisons between mean values of groups were evaluated using an unpaired, 1-tailed Student’s t test. *\*P* ≤ 0.05, *\*\*P* ≤ 0.01 and *** *P* ≤ 0.001.

ID1 inhibits cell senescence in other cell types(*50–52*); therefore, we evaluated senescence in *Tet2^−/−^*chimeric mice (Figure 3I-J). We found that *Tet2^−/−^* LK BMCs showed a significant decrease in the expression of senescence associated marker, β-galactosidase, compared to control mice, which was increased and rescued in *Tet2^−/−^*; *Id1^−/−^* LK progenitors. Taken together, these results suggest that *Id1* intrinsically promotes expansion of *Tet2^−/−^* HSPCs and contributes to CH, in part, by inhibiting apoptosis and senescence of HSPCs.

TET proteins catalyze the conversion of 5-methylcytosine (5mC) to 5-hydroxymethylcytosine (5hmC) that leads to DNA demethylation(*53, 54*). The levels of 5hmC levels are reduced in *Tet2^−/−^*HSPCs and contribute to altered gene expression and multiple phenotypes observed in mice that lack *Tet2^−/−^*. As expected, we found that 5hmC levels are reduced in *Tet2^−/−^*LK cells compared to controls; however, ablation of *Id1* did not rescue 5hmC levels (Figure S5A-B). Thus, *Id1* expression does not affect the levels of 5hmC in *Tet2^−/−^* HSPCs, which could contribute to the hematopoietic phenotypes observed in *Tet2^−/−^* mice.

### *Id1* reduces p16 expression in *Tet2^−/−^* chimeric bone marrow cells that results in reduced apoptosis and senescence

To gain mechanistic insight into how ID1 promotes CH in *Tet2^−/−^*chimeric mice we first evaluated the expression of canonical ID1 targets known to regulate apoptosis, senescence, and proliferation, and then performed bulk RNA sequencing of *Tet2^−/−^*, *Tet2^−/−^*; *Id1^−/−^*and control LK progenitors. ID proteins regulate cell growth, in part, by restraining the transcriptional activity of E and ETS proteins and expression of their downstream targets including cyclin-dependent kinase inhibitors (*Cdki*)(*38, 43, 51, 55–57*); therefore, we evaluated p16, p21, p27 and p57 expression in HSPCs by flow cytometry. The expression of p16 was significantly downregulated in *Tet2^−/−^* donor LK, LSK, Flk2– and MPP cells compared to controls, while p16 expression was increased and rescued in *Tet2^−/−^*; *Id1^−/−^* HSPCs (Frequency and MFI – LK/LSK) (Figure 4A-E).

**Fig. 4.**
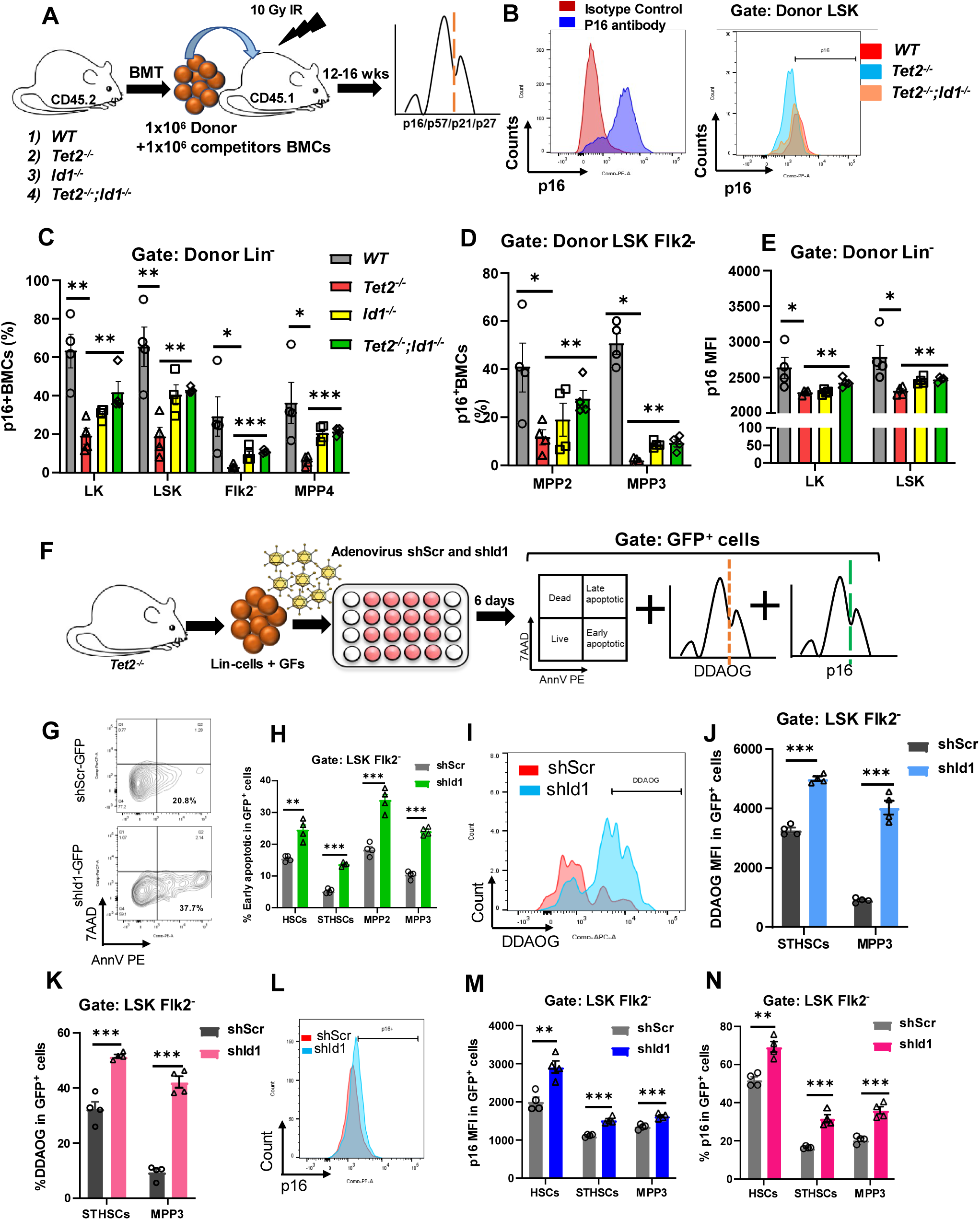
*Id1* inhibits apoptosis and senescence of *Tet2^−/−^* HSPCs by regulating CDKI expression. (A) Summary of experimental design to evaluate cyclin dependent kinase inhibitor (CDKI) expression in donor derived HSPC from competitive BMT recipient mice. (B) Flow cytometry histograms of p16 and isotype control utilized for the gating p16 positive cells. Flow cytometry histograms of p16 expression in donor derived LSK cells in competitive BMT recipient mice. (C) p16 expression in donor derived LK, LSK, Flk2^−^, MPP4, (D) MPP2 and MPP3 in recipient mice by flow cytometry. (E) Mean florescent intensity (MFI) of p16 in donor LK and LSK cells in competitive BMT recipient mice. (F) Summary of experimental procedure to measure apoptosis, senescence and p16 expression in *Tet2^−/−^* Lin^−^ cells transduced with adenoviral vectors expressing scrambled (control) shRNA and GFP (shScr-GFP) and shId1-GFP. (G) Flow cytometry contour plots of *Tet2^−/−^* MPP2 GFP^+^ cells transduced with shScr-GFP and shId1-GFP expressing adenoviral vectors and stained with Annexin V and 7AAD after 6 days of culture. (H) Frequency of early apoptotic GFP*^+^*HSPCs transduced with shScr and shId1 adenoviral vectors after 6 days of culture. (I) Flow cytometry histograms of senescent *Tet2^−/−^* MPP3 (DDAOG^+^) GFP^+^ cells transduced with shScr and shId1 adenoviral vectors. (J) Mean florescent intensity (MFI) and (K) frequency of DDAOG^+^ GFP^+^ cells transduced with shScr and shId1 expressing adenoviral vectors. (L) Flow cytometry histogram of p16 expression in *Tet2^−/−^* MPP3 GFP^+^ cells transduced with shScr-GFP and shId1-GFP expressing adenoviral vectors after 6 days of culture. (M) Mean Florescent Intensity (MFI) and (N) frequency of p16 expression in GFP^+^ cells transduced with *shScr* and *shId1* expressing adenoviral vectors after 6 days of culture. Data are presented as the mean ± SEM, and comparisons between mean values of groups were evaluated using an unpaired, 1-tailed Student’s t test. *\*P* ≤ 0.05, *\*\*P* ≤ 0.01 and *** *P* ≤ 0.001.

Additional analysis of CIP/KIP proteins showed that expression of p57 was decreased in *Tet2^−/−^* LSK, LSK/Flk2^−^, MPP-4, ST-HSCs and MPP3 cells, and was increased and rescued in *Tet2^−/−^*; *Id1^−/−^* LK, LSK/Flk2-and MPP progenitors (Figure S6B-C). Finally, p21 and p27 expression were downregulated in *Tet2^−/−^* HSCs and MPPs compared to control mice and increased and rescued in *Tet2^−/−^* HSCs and MPPs that lack *Id1* (Figure S6D-E). Thus, *Id1* may contribute to the expansion of *Tet2^−/−^* HSPCs and CH by inhibiting p16 and p57 and other CIP/KIP family members.

To support the notion that ID1 promotes the expansion of *Tet2^−/−^*HSPCs by inhibiting p16 expression and reducing apoptosis and senescence, we knocked down (KD) *Id1* expression in *Tet2^−/−^* Lin^−^ cells using adenoviral vectors that express *shId1* and *shScr* (CTL) sequences and GFP, and evaluated, apoptosis, senescence and p16 expression *in vitro* by flow cytometry (Figure 4F). KD of *Id1* in *Tet2^−/−^* Lin^−^ expansion cultures resulted in a significant increase in apoptosis of HSCs, STHSCs, MPP’s after six days (Figure 4G-H) and, increased the MFI and percentage of senescent STHSCs and MPP3’s using a β-gal-sensitive fluorescent probe, DDAOG, (Figure 4I-K). In addition, KD of *Id1* in *Tet2^−/−^* Lin^−^ cells increased the expression (frequency and MFI) of p16 in HSCs, STHSCs and MPP3 by flow cytometry (Figure 4L-N). Thus, reducing *Id1* expression in *Tet2^−/−^* HSPCs BMCs increases p16 expression, apoptosis, and senescence of HSPCs.

To establish that p16 expression regulates apoptosis and senescence of *Tet2^−/−^* HSPCs, we knocked down *p16* expression in *Tet2^−/−^*; *Id1^−/−^* Lin^−^ cells using lentiviral vectors that express *shp16* or *shScr* (Figure 5A). The expression of p16 was significantly reduced in p16 KD Lin-cells (Figure 5B), which resulted in a significant decrease in the percentage of early apoptotic Lin-cells (Figure 5C), and C_12_FDG^+^ senescent STHSCs and MPP3 cells by flow cytometry (Figure 5D). Taken together, p16 promotes apoptosis and senescence of *Tet2^−/−^*; *Id1^−/−^* Lin^−^ cells and suggest that ID1 inhibits apoptosis and senescence of *Tet2^−/−^*cells by inhibiting p16 expression.

**Fig. 5.**
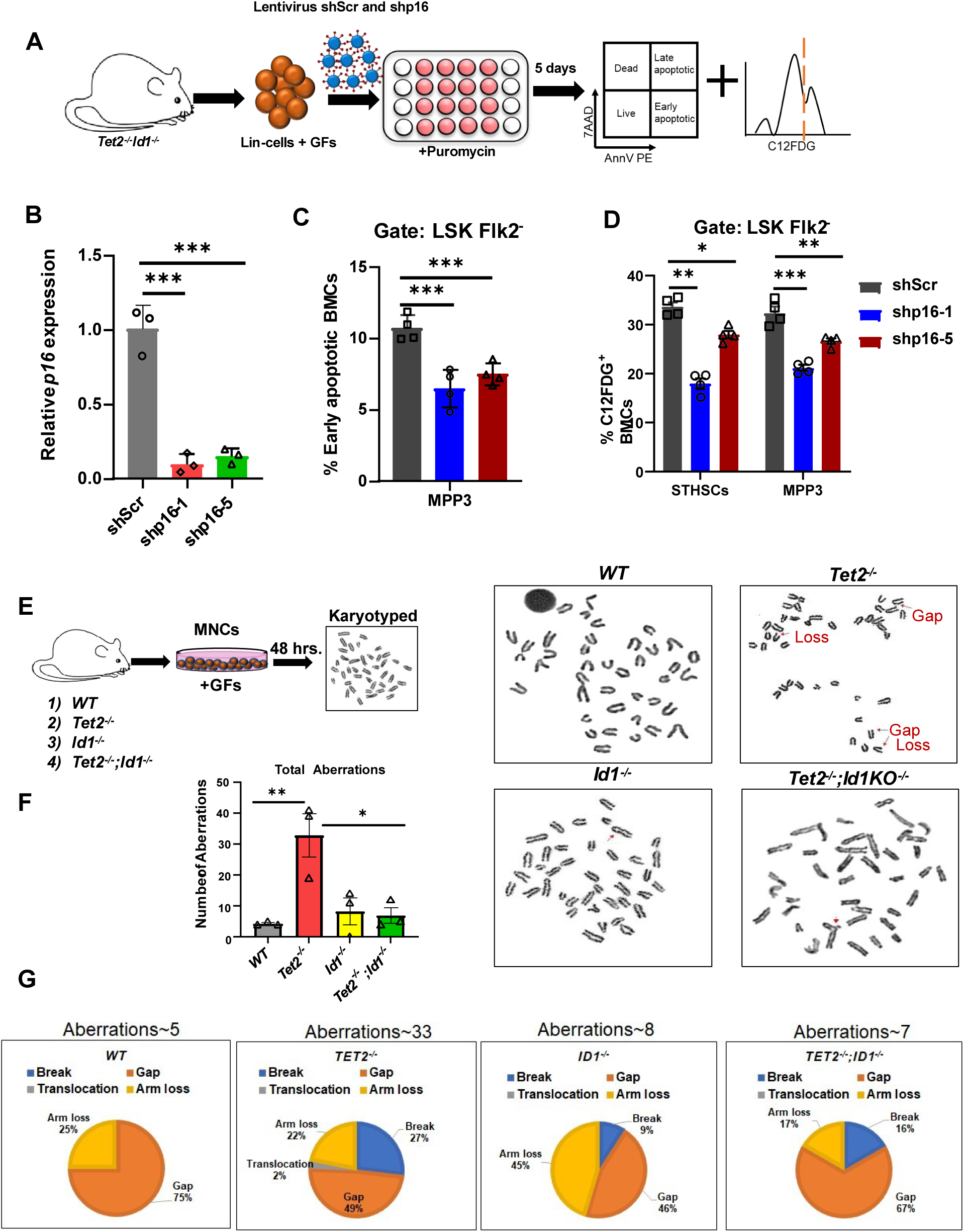
p16 regulates apoptosis and senescence of *Tet2^−/−^*HSPCs and ablation of Id1 rescues chromosomal instability of *Tet2^−/−^*BMCs. (A) Summary of experimental design for lentivirus shRNA mediated knockdown of p16 expression in *Tet2^−/−^;Id1^−/−^*Lin^−^ cells to assess apoptosis and senescence. (B) RT-qPCR analysis of p16 expression in the cultured *Tet2^−/−^;Id1^−/−^*cells, in the presence of puromycin, after knockdown of p16 expression. (C) Percentage of early apoptotic MPP3 cells in shScr and shRNA p16 clone#1 (shp16-1) and shRNA p16 clone#5 (shp16-5) after 5 days of culture. (D) Frequency of senescent C_12_FDG^+^ STHSCs and MPP3 cells in shScr, shp16-1 and shp16-5 transduced *Tet2^−/−^; Id1^−/−^*Lin^−^ cells after 5 days of culture. (E) Summary of experimental procedure for karyotype analysis of *Tet2^−/−^*, *Tet2^−/−^;Id1^−/−^* and control mononuclear cells cultured in GFs for 48 hrs. (F) Photomicrographs of chromosome spreads and quantitation of chromosomal aberrations observed in metaphase spreads of *Tet2^−/−^, Tet2^−/−^;Id1^−/−^* and control mononuclear cells after culturing in GFs for 48 hrs. (G) Venn-diagram showing types of chromosomal aberrations present in cultures of indicated cells. Data are presented as the mean ± SEM, and comparisons between mean values of groups were evaluated using an unpaired, 1-tailed Student’s t test. *\*P* ≤ 0.05, *\*\*P* ≤ 0.01 and *** *P* ≤ 0.001.

To gain further mechanistic insights into how ID1 promotes CH in *Tet2^−/−^* mice, we analyzed differentially expressed genes in LK RNA-seq data sets by Ingenuity Pathway and GSEA analysis (Table S1, Figure S6F and Figure S7A-B). As predicted, ID1 signaling pathway genes were upregulated in *Tet2^−/−^* compared to *Tet2^−/−^*; *Id1^−/−^* LK cells. ERK/MAPK, IL-8/IL-1/IL-6, chemokine and mTOR signaling pathway genes were upregulated in *Tet2^−/−^* LK compared to *Tet2^−/−^*; *Id1^−/−^* LK cells indicating that the *Tet2^−/−^* LK cells were exposed to an inflammatory environment that was rescued by loss of *Id1*, which is likely due to a reduction in the number of proinflammatory cells. p53 pathway and apoptosis pathway genes, and cell cycle checkpoint/G2/M phase/kinetochore pathway genes were decreased in *Tet2^−/−^* LK cells compared to *Tet2^−/−^*; *Id1^−/−^* LKs cells, which could contribute to decreased genomic stability in *Tet2^−/−^*mice. Finally, IPA analysis showed that RHO GTPase cycle genes were significantly upregulated in *Tet2^−/−^* LK compared to *Tet2^−/−^*; *Id1^−/−^* LK cells, which suggest an unexplored mechanism of Id1 action in promoting cell survival, proliferation, and self-renewal(*58*). GSEA analysis confirmed the IPA analysis and found that *Tet2^−/−^*; *Id1^−/−^*LK cells show 1) decreased inflammatory gene expression, 2) decreased, and TOR signaling pathway genes, and 3) increased DNA repair, chromosome separation, and G2M checkpoint gene expression compared to *Tet2^−/−^*LK cells (Figure S8A-D). Taken together, IPA and GSEA pathway analysis confirms that 1) inflammatory signaling pathways are downregulated in *Tet2^−/−^*; *Id1^−/−^*LK cells, 2) that Id1 may regulate the p53 pathway and contribute to the decreased apoptosis and genomic instability of *Tet2^−/−^* LK cells, and 3) Id1 may promote CH by upregulating RHO GTPase signaling and function.

### Ablation of *Id1* rescues chromosomal instability of *Tet2^−/−^* BMCs

TET proteins regulate DNA damage response pathway protein expression, and cells with loss of TET function show increased DNA strand breaks, genomic instability, and develop myeloid malignancies(*20, 59–62*). Therefore, we cultured *Tet2^−/−^* and *Tet2^−/−^*; *Id1^−/−^* BMCs in HGFs *in vitro* and analyzed mitotic cells for chromosomal aberrations (Figure 5E). There was a significant increase in chromosomes with gaps, breaks, arm loss and translocations in *Tet2^−/−^* BMCs compared to controls (Figure 5F-G). Chromosomal aberrations were reduced and rescued in *Tet2^−/−^*; *Id1^−/−^*BMCs in three separate experiments demonstrating that ablation of *Id1* reduces genomic instability of *Tet2^−/−^* BMCs. Since TET1 regulates the expression of DNA repair genes including BRCA1, RAD51 and 53BP1, and loss of TET1/2 leads to genomic instability, we evaluated RAD51 and 53BP1 expression in *Tet2^−/−^*chimeric mice by flow cytometry (Figure S9A). The expression of RAD51 and 53BP1were significantly reduced in *Tet2^−/−^*HSPCs compared to controls and were increased and rescued in *Tet2^−/−^*; *Id1^−/−^* HSPC (Figure S9B-E). Thus, loss of *Id1* expression results in increased DNA repair protein expression that is correlated with increased genomic stability in *Tet2^−/−^* HSPCs.

Since *Tet2^−/−^* BMCs show reduced apoptosis (Figure 3A-B), increased genomic instability (Figure 5E-G), reduced expression of p53 pathway signaling genes (Figure S7B) and, ID1 down regulates p53 expression in other cell types(*63, 64*); we evaluated p53 and p19 expression in HSPCs from chimeric mice (Figure 6A). The expression of p53 was reduced in *Tet2^−/−^* LK, LSK, ST-HSC and MPP cells compared to control cells (Figure 6B and Figure S9F-G); however, p53 expression was selectively increased and rescued in *Tet2^−^; Id1^−/−^* LK cells but not LSK cells. The expression of p19, a positive regulator of p53, was similarly reduced in LK, LSK, and MPP, and rescued in LK cells (Figure 6C and Figure S9H). Taken together, p53 expression is downregulated in primitive and committed *Tet2^−/−^* HSPCs and rescued by loss of *Id1* function in committed *Tet2^−/−^* progenitors, suggesting that p53 may function to increase genomic stability in more committed progenitors.

**Fig. 6.**
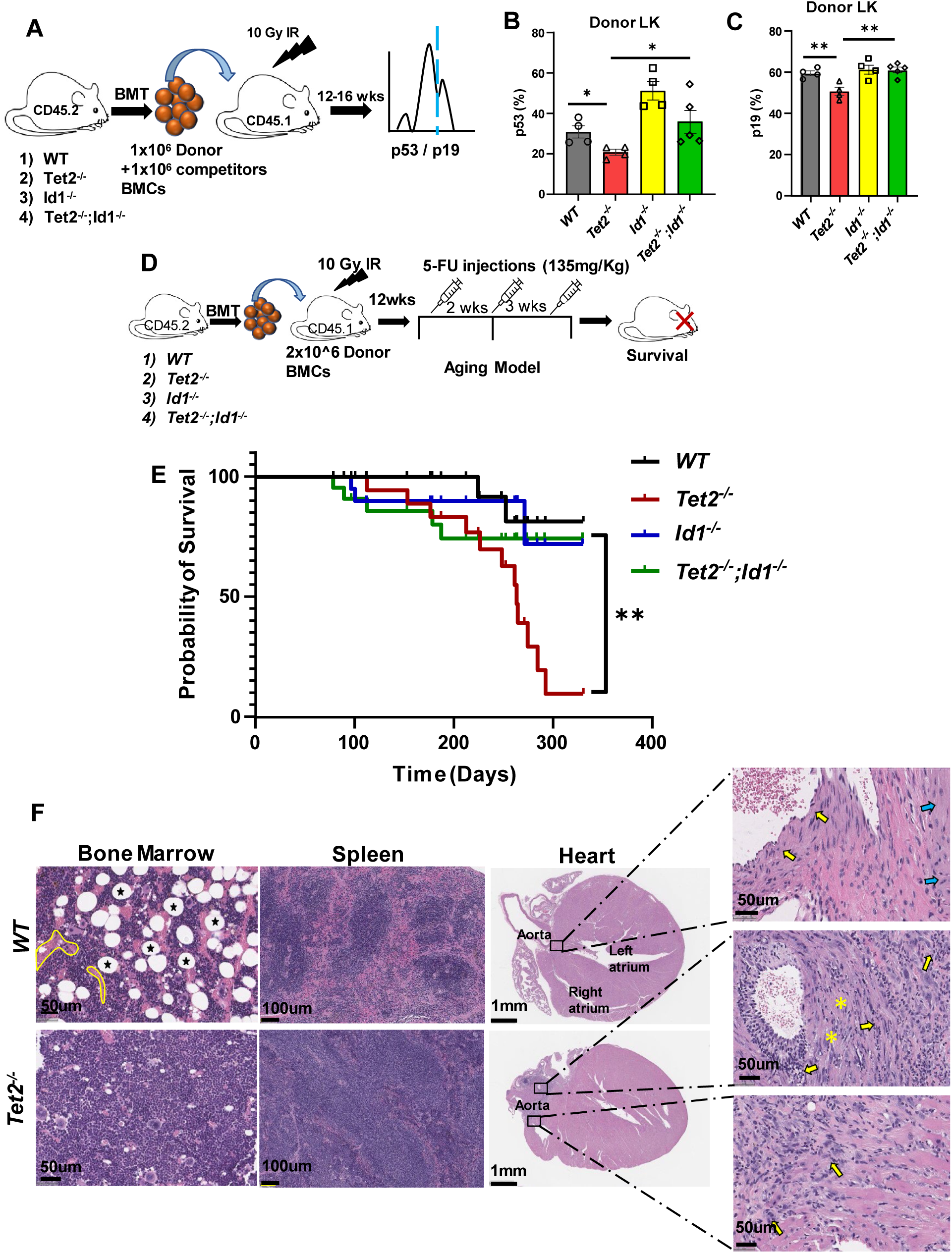
*Id1* ablation delays the onset of disease progression in *Tet2^−/−^* chimeric mice. (A) Summary of experimental procedure to evaluate p53 and p19 expression in donor derived HSPC from competitive BMT recipient mice. (B) Percentage of donor derived LK cells that express p53 in BMT recipient mice. (C) Percentage of donor derived LK cells that express p19 in BMT recipient mice. (D) Summary of experimental design to treat noncompetitive BMT recipient mice with 5-FU to accelerate aging. (E) Kaplan-Meier survival curve of 5-FU treated noncompetitive BMT recipient mice. (F) Representative hematoxylin and eosin-stained sections of BM, spleen and heart from moribund mice transplanted with *Tet2^−/−^* cells and mice transplanted with *WT* cells. *WT* BM sections show normal hematopoietic cell, adipocytes (black stars), and vascular lumen (yellow) distribution, while *Tet2^−/−^* BM sections show densely packed hypercellular BM with reduced adipocyte and fibrovascular tissue. WT spleens showed normal red and white pulp (yellow outline), while *Tet2^−/−^* spleens showed hematopoietic cell hyperplasia and disrupted splenic architecture (yellow outline). WT heart sections show normal myocardium (blue arrow) and aorta (yellow arrow). *Tet2^−/−^* heart sections show increased numbers of inflammatory cells observed within the vessel wall and surrounding vessels (yellow arrow), and loss and fragmentation of elastic fiber (yellow asterisk). Scale bar, 50um for bone marrow, 100um for spleen, 1mm for heart and 50um for magnified heart section. Data are presented as the mean ± SEM, and comparisons between mean values of groups were evaluated using an unpaired, 1-tailed Student’s t test. Kaplan-Meier survival studies were analyzed using Wilcoxon’s signed-rank. *\*P* ≤ 0.05, *\*\*P* ≤ 0.01.

### *Id1* ablation delays the onset of disease progression in *Tet2^−/−^* chimeric mice

Ablation of *Id1* in *Tet2^−/−^* chimeric mice reduces 1) HSPC expansion/self-renewal, 2) extramedullary hematopoiesis, 4) myeloid skewing, and 5) chromosomal abnormalities; therefore, we reasoned that loss of *Id1* may delay or reduce the onset or incidence of leukemia. *Tet2^−/−^* mice develop myeloid and lymphoid hyperplasia that leads to hematopoietic malignancies in some mice after a long latency(*17, 19, 65–67*); therefore, we treated chimeric mice with multiple sublethal doses of 5-FU to accelerate aging, by promoting proliferative stress and accumulation of secondary mutations and disease (Figure 6D)(*68*). *Tet2^−/−^* chimeric mice became sick and moribund beginning 100 days after transplantation and treatment with 5-FU, while control chimeric mice survived (Figure 6E). Histopathological examination of moribund *Tet2^−/−^* chimeric mice showed BM hyperplasia with densely packed BM, and splenic hyperplasia that altered the normal architecture of the spleen (Figure 6F). Increased cardiac inflammation/vasculitis was observed in some sick *Tet2^−/−^* chimeric mice compared to aged matched healthy *Tet2^+/+^* mice, which is likely due to increased cardiac inflammation caused by γ-IR conditioning for BMT and made worse by hyperinflammatory *Tet2^−/−^* macrophages (Figure 6F)(*21, 26, 47*). Thus, loss of *Id1* in *Tet2^−/−^* chimeric mice reduces hematopoietic hyperplasia and inflammation and delays the onset of leukemia.

### The small molecule inhibitor of ID1, AGX51, inhibits the expansion of *Tet2^−/−^* chimeric HSPCs *in vitro*

Since loss of ID1 function delays the onset of disease in *Tet2^−/−^* chimeric mice we speculated that the recently identified small molecule inhibitor of ID1, AGX51, could reduce HSPC expansion in this mouse model(*69*). AGX51 significantly inhibited the enhanced colony forming ability of *Tet2^−/−^* BMCs (Figure 7A). The inhibitory effects of AGX51 on WT BMC colony formation was expected since this is a stress assay promoted by growth factors that induce *Id1* including IL-3(*39, 40, 43*). Similarly, AGX51 inhibited the proliferation and growth of WT and *Tet2^−/−^* MPP cells in BrdU-incorporation (Figure 7B-C) and cell viability assays (Figure 7D-E). Collectively, AGX51 inhibits the expansion of *Tet2^−/−^* HSPCs *in vitro* and may represent a potential therapy to treat CH and delay onset and progression of leukemia.

**Figure 7.**
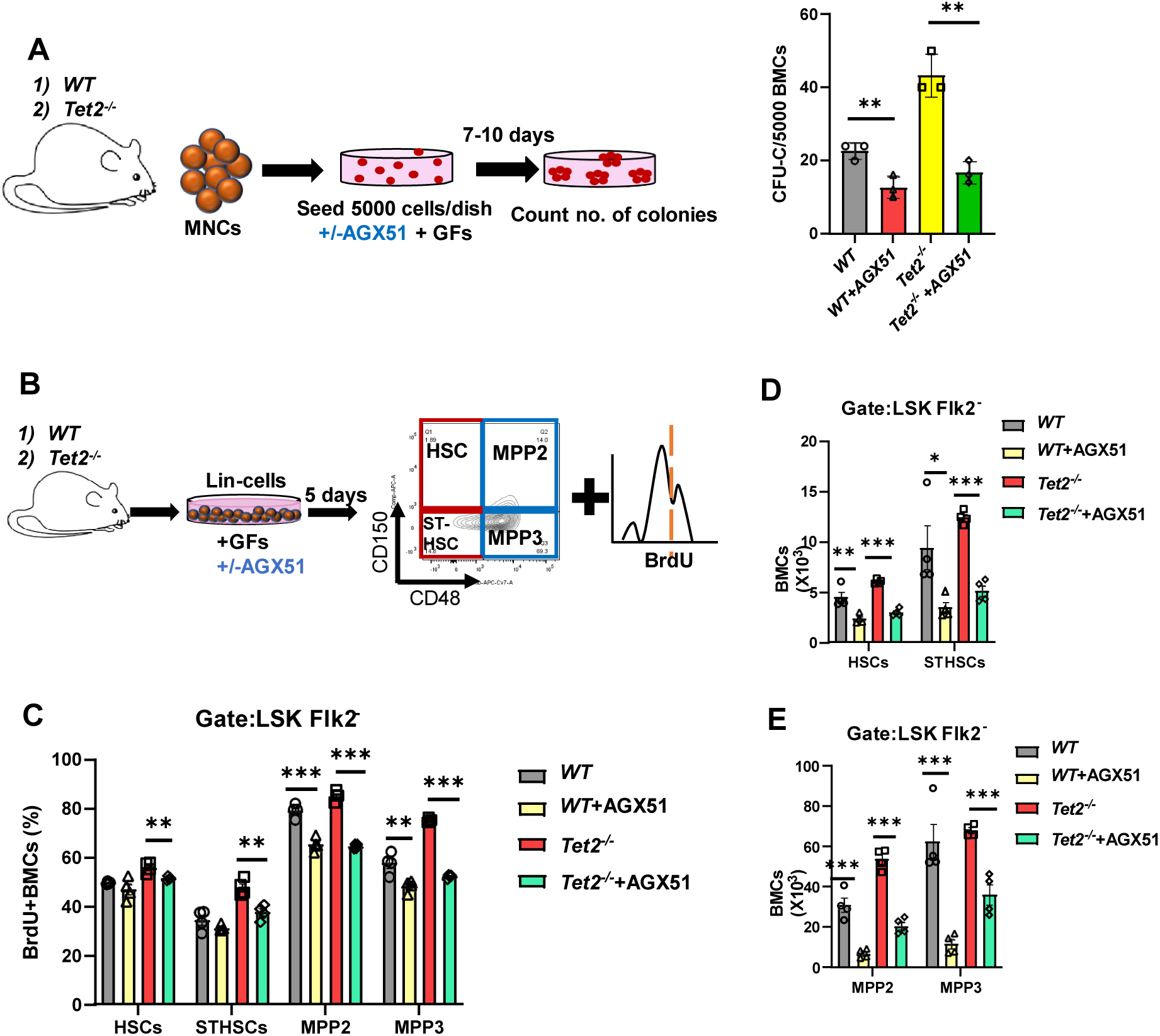
AGX51 inhibits the expansion of *Tet2^−/−^* BM HSPCs. (A) Summary of experimental procedure to measure the number of colony forming progenitor cells (CFU-c) in *WT* and *Tet2^−/−^*BMCs, and the number of CFU-c in 5 x 10^3^ AGX51 treated and control BMCs. (B) Summary of experimental procedure to evaluate expansion and proliferation potential of the *Tet2^−/−^* BMCs treated with AGX51. (C) Percentage of BrdU^+^ *Tet2^−/−^* HSPCs in Lin^−^ cell cultures treated with AGX51 after 5 days. (D) Total number of HSCs and STHSCs and (E) MPP2 and MPP3 in *Tet2^−/−^* Lin^−^ cells cultured in the presence or absence of AGX51 after 5 days. Data are presented as the mean ± SEM, and comparisons between mean values of groups were evaluated using an unpaired, 1-tailed Student’s t test. *\*P* ≤ 0.05, *\*\*P* ≤ 0.01 and *** *P* ≤ 0.001.

## DISCUSSION

Chronic inflammation promotes HSPC expansion and contributes to CH, MPN and AML(*31, 33, 70–72*). Cytokines and pro-inflammatory mediators produced during chronic inflammatory stress induce *Id1* in HSPCs and promote proliferation, suggesting that ID1 could contribute to CH (*39, 40, 43, 73*). We show that genetic ablation of *Id1* in a mouse model of CH reduces HSPC expansion and self-renewal, myeloid skewing, extramedullary hematopoiesis, genetic instability, and disease onset, which is correlated with increased differentiation, apoptosis and senescence, and decreased proliferation (*51, 52*). Mechanistically, *Tet2^−/−^* HSPCs show reduced p16/CDKI expression, and loss of *Id1* expression in *Tet2^−/−^*HSPCs *in vivo* and *in vitro* increases p16/CDKI expression, and promotes senescence and apoptosis, and reduces cell growth, which collectively contribute, in part, to reducing CH. We speculate that ID1 increases p16/CDKI expression via the canonical pathway of antagonizing E protein transcriptional activity (ID-E-CDKI pathway)(*51, 55–57, 74*). We provide evidence that a small molecule inhibitor of ID1, AGX51, reduces the expansion of Tet*2^−/−^* HSPCs *in vitro* suggesting that AGX51 could be employed as a first line defense to inhibit the expansion of preleukemic clones in patients with CH with high-risk progression to AML.

Inhibition of inflammation, inflammatory cytokines and downstream intracellular signaling pathways are potential therapeutic targets to reduce CH (*23, 24, 31–33, 71, 75*). Administration of neutralizing IL-6 antibodies and inhibitors of signaling proteins downstream of IL-6 suppress CH in *Tet2^−/−^*mice(*23, 25*). Ablation of *IL-1r1* expression in *Tet2^−/−^* HSPCs rescues LSK expansion, myeloid skewing, and the pro-inflammatory state(*29*). Thus, inhibiting inflammatory signaling pathways in patients with *TET2* mutations and CH could reduce HSPC expansion and delay progression to MPN and hematopoietic malignancies. However, cytokines can act alone or in combination to promote CH; therefore, it is difficult to predict which cytokines or cytokine combinations to inhibit, and some patients might show increased risk of infection after blocking cytokine function. Since *Id1* is expressed at low levels in primitive HSPCs during steady state hematopoiesis and is induced after BMT, inflammatory and genotoxic stress by pro-inflammatory cytokines (*39, 40, 43, 73*), inhibitors of ID1 represent attractive alternative therapies to reduce CH. Thus, it will be important to extend our initial findings that AGX51 reduces *Tet2^−/−^*HSPCs expansion *in vitro* to inhibition of CH in *Tet2^−/−^*mice *in vivo* and human models of CH, MPN, AML and CMML *in vitro and vivo*, and evaluate other small molecule inhibitors of ID1 including cannabidiol (*76, 77*). ID1 inhibitors might be therapeutic in patients with other mutations that promote CH including JAK2, or more broadly in patients with MPN with chronic inflammation and MDS, since *Id1* is induced downstream of JAK2 signaling and other pro-inflammatory signals(*73, 78*). Intriguingly, ID1 is frequently overexpressed in many cancers, and clonal expansion and chronic inflammation is observed in other tissues during aging and tumor progression suggesting that inhibiting ID1 may slow early-stage progression of other cancers(*79–83*).

Current studies support the hypothesis that *Tet2^−/−^*immune cells are hyperinflammatory and produce elevated levels of proinflammatory cytokines that indirectly promote HSPC expansion and contribute to CH. However, loss of TET2 function could intrinsically regulate Id1 gene expression; therefore, future studies will examine if loss of *TET2* function intrinsically regulates *Id1* gene methylation and expression in HSPCs shortly after *Tet2* gene deletion *in vitro and in vivo*.

Patients with *TET2* mutations and CH have an increased risk of developing atherosclerosis and cardiovascular disease (CVD)(*84–86*). Atherosclerosis prone *Ldlr^−/−^* mice transplanted with *Tet2^−/−^*BMCs show increased inflammation and hyperinflammatory macrophages that promote atherosclerotic plaque formation(*4, 21, 84*). *Tet2^−/−^*; *Id1^−/−^* chimeric mice show reduced myeloid cell expansion suggesting that genetic ablation of *Id1* in *Tet2^−/−^;Ldlr^−/−^*mice may reduce and rescue atherosclerosis(*22, 87*). Thus, inhibiting *Id1* expression or function in patients with *TET2* mutations and CH might reduce the risk of atherosclerosis and CVD.

*Tet2^−/−^* HSPCs show increased genomic instability that is correlated with a decrease in the expression of DNA repair proteins(*20, 59, 61*). Unexpectedly, we found that ablation of *Id1* expression in *Tet2^−/−^* HSPCs increases genomic stability that was correlated with 1) an increase in DNA repair protein gene expression, RAD51 and 53BP1, 2) increased p53 protein expression (LK cells), 3) increased G2M checkpoint pathway gene expression, and increased sister chromatin separation gene expression, and 4) decreased proliferation, which could contribute to increased genomic stability. Mechanistically, *Id1* has been reported to inhibit p53 expression; however, the precise mechanism(s) that ID1 regulates DNA repair protein expression in HSPCs and p53 expression in LK cells is currently not known(*63, 64*). Additional experiments are needed to uncover if these, or other proteins and pathways not yet identified, mediate the increase in genomic stability observed in *Tet2^−/−^*; *Id1^−/−^*HSPCs.

In summary, we provide evidence that genetic ablation of *Id1* in *Tet2-*deficient HSPCs rescues hematopoietic phenotypes associated with CH in these mice. Furthermore, mice that lack *Id1* develop and age normally suggesting that specific inhibitors of ID1 may be therapeutic and well tolerated in patients with *TET2* mutations or other mutations that promote inflammation and CH (*88*).

## MATERIALS AND METHODS

### Mice

Id1^−/−^ mice on C57BL/6 background were derived from Id1^−/−^ mixed background mice (B6;129 generously provided by Robert Benezra) after 10 (or greater) generations of backcross with C57BL/6 mice (Charles River). Tet2^−/−^ mice were purchased from Jackson laboratory (023359). Id1^−/−^ mice were bred with Tet2^−/−^ mice to generate *Tet2^−/−^;Id1^−/−^*mice. C57BL/6 Id1-EGFP knock-in reporter mice (*Id1GFP/+*) were generously provided by Xiao-Hong Sun, Oklahoma Medical Research Foundation, Oklahoma, US. C57BL/6 Ly5.2 (CD45.1) mice were obtained from Charles River. All mice were maintained on C57BL/6 genetic background.

Recipient mice used in BMT experiments were 8–12-week-old females. [Experiments involving the use of mice were approved by the NCI at Frederick Animal Care and Use Committee in accordance with the eighth edition ‘‘Guide for the Care and Use of Laboratory Animals.’’]

### Cell lines

HEK 293T/17 cells were cultured in 10% Fetal bovine serum (Gibco, 15140) 1% penicillin/streptomycin in Dulbecco’s modified eagle medium. Transfection studies in HEK 293T/17 cells were performed following the guidelines outlined for LipoD293™ In Vitro DNA Transfection Reagent (Signagen, SL100668) in 100mm dish. Mo7e (DSMV# ACC 104), K562 (ATCC# CRL-3343), KG1a (ATCC# CCL-246.1) cells were grown as described by the supplier.

### Bone marrow transplantation

For competitive bone marrow transplantation assays, 1×10^6^ BMCs from WT (*Tet2^+/+^; Id1^+/+^*), *Tet2^−/−^* (*Tet2^−/−^; Id1^+/+^*)*, Id1^−/−^* (*Tet2^+/+^; Id1^−/−^*), and *dKO* (*Tet2^−/−^; Id1^−/−^*) mice (CD45.2) were mixed with 1×10^6^ C57BL/6 (CD45.1) BMCs and transplanted into γ-irradiated (10 Gy, 137Cs source) congenic CD45.1 mice by tail vein injection (iv). Recipient mice were pretreated one week before and two weeks after transplantation with antibiotic-containing water (pH 2.5-3.0, 0.5 mg/mL amoxicillin, 0.17 mg/mL enrofloxacin). Donor reconstitution was assessed by flow cytometry in the bone marrow and/or peripheral blood after 12-16 weeks for all BMT recipient mice.

### Flow cytometry analysis of hematopoietic stem and progenitor cells

Single-cell suspensions were prepared from *WT, Tet2^−/−^, Id1^−/−^* or *dKO* BMCs or from BMCs of animals transplanted with them. BMCs were harvested from femurs and tibias by flushing with 1% BSA PBS using 25-gauge needle and syringe filtered through a 40 μm membrane. The light density BMC fraction was isolated using lymphocyte separation media (LSM) (MO Biomedicals).

Peripheral Blood Cells (PBCs) collected from mandibular bleed was stained with cell surface markers and incubated with BD Pharm lysis buffer for 10 min., to remove RBCs and washed in 1% BSA PBS solution, before acquiring. For lineage analysis, PBCs and BMCs were incubated with the following fluorochrome conjugated anti-mouse monoclonal antibodies: CD45.2 (104), CD11b (M1/70), Gr-1 (RB6-8C5), CD19 (1D3), CD11c (HL3), F4/80 (AG BM8), Ly6C (AL-21), Ly6G (1A8), CD4 (GK1.5), CD8 (53-6.7), CD71(R17217) and Ter119 (Ter119). For HSPC analysis, LSM purified BMCs were treated with Fc receptor blocking antibodies (anti-mouse CD16/32), then incubated with fluorochrome-conjugated mouse lineage antibody cocktail from BD, c-Kit (ACK2), Sca-1(D7), CD150 (mShad150), CD48 (HM48-1), CD34 (RAM34), Flk2 (A2F10), anti-FcgRII/III (FcR) (93). Lin-negative (Lin^−^) c-Kit^+^ and Sca-1^+^ (LSK) cells were analyzed according to the following marker expression: HSC: LSK Flk2^−^CD150^+^ CD48^−^, short-term HSC (STHSCs): LSK Flk2^−^CD150^−^CD48^−^, multipotent progenitors (MPP2): LSK Flk2^−^ CD150^+^CD48^+^ and multipotent progenitors (MPP3): LSK Flk2^−^CD150^−^CD48^+^. All cells were incubated in staining buffer (1%BSA PBS) for 45 min on ice with intermittent mixing, and then washed in buffer prior to acquiring. All the antibodies used were purchased from BD Biosciences (San Jose, CA) or eBiosciences (San Diego, CA).

### *Id1-GFP* reporter cell assays

For the evaluation of *Id1* expression in the *Tet2^−/−^*mouse model, *Id1GFP/+* mice were bred with *Tet2^−/−^* mice resulting in *Tet2^−/−^; Id1GFP/+* mouse model. BMCs were harvested from *Id1GFP/+* and *Tet2^−/−^; Id1^GFP/+^* mice and fractionated using LSM. The cells were then stained for HSCs, ST-HSCs, MPPs, common myeloid progenitors (CMP), Lin-c-Kit^+^ (LK) CD34^+^ FcR^−^, granulocyte/macrophage progenitors (GMP), LK CD34^+^FcR^+^, and megakaryocyte erythroid progenitors (MEP) LK CD34^−^FcR^−^. The cells were fixed and permeabilized with BD cytofix/perm buffer, and then blocked with 5% BSA in PBS for 45 min. on ice and then stained with primary antibody, chicken (chk) anti-GFP (ab 13970) for 1 hr. The cells were then washed and incubated with secondary antibody, anti-chk AF488 (ab 150169) for 30 min., washed twice with PBS and analyzed for GFP expression by flow cytometry.

### Intracellular staining

For analysis of cyclin dependent kinase inhibitor (CDKI) and DNA damage response protein expression, all antibodies were purchased from Santacruz Biotech. BMCs stained for HSPC analysis were fixed and permeabilized with BD cytofix/perm buffer, and then blocked with respective purified isotype control antibody for 40 min on ice and then stained with p16 (sc-166760 PE), p21 (sc-6246 AF488), p27 (sc-1641PE), p57 (sc-56341 AF488), p53 (sc-126 PE), p19 (sc-32748 AF488), Rad51 (sc-398587 AF594) and 53BP1 (sc-515841 AF488) for 45 min. on ice and analyzed by flow cytometry.

### Survival assay

Chimeric mice were generated by transplanting 2×10^6^ BMCs from WT, *Tet2^−/−^, Id1^−/−^*, and *Tet2^−/−^;Id1^−/−^* mice (CD45.2) into γ-irradiated (10 Gy, 137Cs source) recipient mice by tail vein injection (iv). Twelve weeks after BMT, recipient mice were treated with three doses of 5-fluorouracil (5-FU) (135mg/kg) via intraperitoneal injections, at an interval of 2 weeks and 3 weeks, respectively and monitored for survival. Sick mice were euthanized in accordance with the “Guide for the Care and Use of Laboratory Animals.”

### Proliferation and Cell Cycle analysis

Chimeric mice were generated by transplanting 2×10^6^ (1:1 Donor+Competitor) *WT, Tet2^−/−^, Id1^−/−^* and *Tet2^−/−^; Id1^−/−^* BM cells into 10 Gy irradiated recipient mice. Twelve weeks post transplantation the recipient mice were subjected to one intraperitoneal BrdU injection (1mg/6 gm mice) (Sigma, B5002). These mice also received BrdU (1mg/ml BrdU+1 %Glucose) in drinking water for 2 days. The BMCs were harvested and stained for HSPC markers, fixed, permeabilized and re-fixed with Cytofix/Cytoperm buffer (BD Biosciences) followed by DNase I (Sigma, D4513) treatment (300ug/ml) for 1 hr at 37°C. The cells were washed and stained with anti-BrdU antibody (BD Biosciences) for 30min. at RT, washed and analyzed by flow cytometry. For cell cycle/quiescence intracellular staining, BMCs stained for HSPC analysis were fixed and permeabilized with Cytofix/Cytoperm buffer (BD Biosciences) followed by 40 min. intracellular staining with Ki-67-PE (B56) (BD Biosciences, 567719). The cells were then incubated overnight with FxCycle Violet dye (ThermoFisher, F10347) and acquired next day. BD LSRII SORP, BD Fortessa or BD FACSymphony A5 were used for FACS acquisition and Aria II for cell sorting. Data analysis was done using FlowJo V9 software (Tree Star Inc., Ashland OR).

### Apoptosis Assays

Chimeric mice were generated by transplanting 2×10^6^ (1:1 Donor + Competitor) *WT, Tet2^−/−^, Id1^−/−^* and *Tet2^−/−^; Id1^−/−^* BM cells into 10 Gy irradiated recipient mice. 12 weeks post transplantation the BMCs were harvested and stained for HSPC markers, fixed, permeabilized, labeled with TUNEL reaction mixture, as per the protocol (Sigma, #11684795910). The cells were washed twice and analyzed by flow cytometry. For annexin V staining, BMCs were incubated with annexin V-PE and 7AAD (BD Biosciences 559763) for 15 min. at RT and further stained for HSPC analysis and immediately acquired on flow cytometer.

### Senescence assay

*In vitro* cultured cells or BMCs from transplanted recipient mice were harvested and stained for HSPC markers. The cells were then resuspended in growth media and treated with a final concentration of 100nM of bafilomycin A1(Sigma, B1793) for 1 hr at 37°C, C_12_FDG (Thermofisher Scientific, #D2893) at a final concentration of 33uM, or DDAOG (Thermofisher Scientific, #D6488) at a final concentration of 20uM was added on to the cells and incubated for 1 hr at 37°C. The cells were further stained with the HSPC markers and analyzed by flow cytometry.

### Hematopoietic Stem and Progenitor Expansion Assays. *In vitro* studies

LSM BMCs were isolated from experimental mice and incubated with biotinylated Mac-1 (M1/70), Gr-1 (RB6-8C5), B220 (RA3-6B2), CD4 (GK1.5), CD8 (53-6.7), IL7R (A7R34) and Ter119 (Ter119) and Lin+ IL7R+ cells were subsequently removed with streptavidin beads. Lineage negative, Lin-cells were then cultured in StemSpan (#09650) medium containing mSCF (Peprotech# 250-03) (100 ng/mL), hTPO (Peprotech# 300-18) (100 ng/mL), mFGF1 (Peprotech# 450-33A) (10ng/mL), hIGFII (Peprotech# 100-12) (20 ng/mL), and Angiopoietin2 (Peprotech# 130-07) (50ng/mL). Cells were removed from culture after 5-6 days and stained with HSPCs markers. To monitor proliferation, using BrdU, cultured Lin^−^ cells were pulsed with 1mM BrdU (BD, #559619) 30-40 min. before harvest, stained for HSPCs, fixed and permeabilized with Cytofix/Cytoperm buffer (BD Biosciences) followed by 40 min. intracellular staining with anti-BrdU antibody and analyzed by flow cytometry.

### *In vitro* differentiation culture

LSM BMCs were isolated from *WT, Tet2^−/−^, Id1^−/−^*and *Tet2^−/−^; Id1^−/−^* mice. Lin^−^ cells were cultured in IMDM+10% FBS containing mSCF (50ng/ml), mFlt3 (10ng/ml), mIL3(10ng/ml), MCSF (10ng/ml), GCSF (10ng/ml), GMCSF (10ng/ml). After 6 days of culture the cells were harvested, stained with lineage specific antibodies, and analyzed on flow cytometer.

### Colony formation assay

BMCs were harvested from *WT, Tet2^−/−^, Id1^−/−^* and *Tet2^−/−^; Id1^−/−^* mice and labeled with antibodies that recognize HSPC as described above, and HSCs (LSK/ FLT3^−^/CD150^+^/CD48^−^) were FACS sorted HSCs per plate were seeded in methocult (Stemcell technologies, M3434) containing 35mm gridded dishes. Colonies were scored 7-10 days after the seeding, under the microscope. For serial replating, colonies in a single plate were harvested together, counted and 5000 cells/plate were reseeded in the methocult for screening after another 7-10 days. Up to 7 replating were performed for the experiments.

### Single cell Division Kinetics assay

BMCs were harvested from *WT, Tet2^−/−^, Id1^−/−^*and *Tet2^−/−^; Id1^−/−^* mice and labeled with antibodies that recognize HSPC as described above. LK (Lin^−^ ckit^+^) and HSCs (LSK/ FLT3^−^/CD150^+^/CD48^−^) were sorted and single cell deposited directly into

Terasaki plates (Greiner Bio-One) containing StemSpan serum free media supplemented with mSCF (100ng/ml), hTPO (100ng/ml), IL-6 (50ng/ml) and mIL-3 (30ng/mL) in vitro. Wells with percent confluency were counted at 7 days after seeding.

## Acknowledgements

We wish to thank Lai Thang, Christina Robinson and the LASP Animal Research and Technical Support Staff for providing technical assistance in animal studies. Jeff Carrell and Megan Karwan in the NCI-Frederick Flow Cytometry Core for cell sorting. Bao Tran and Keyur Talsania in CCR-Sequencing facility for RNA-Sequencing. Melissa Galloux for RNA-Sequencing data analysis. Jennifer Matta in the FNLCR-Molecular Histopathology laboratory for the sick mice organ section staining and scanning. Stephen Fox from Protein Characterization FNLCR Core facility for LC-MS. This study makes use of data generated by Department of Pathology and Tumor Biology, Kyoto University(*89*).

## Author Contributions

Conceptualization: SS, TS, BLJ, KOG, JRK

Conducted experiments: SS, TS, BLJ, SB, KOG, HMM, GTP, DMS, JPS, JRK

Analyzed Results: SS, TS, BLJ, KOG, SB, BK, JRK

Writing – original draft: SS, JRK

Writing – review and editing: SS, JRK, KOG, TS

