## Supplemental Figures – methods for "*Id1* Promotes Clonal Hematopoiesis in Mice with *Tet2* Loss of Function"

Shweta Singh *et al.*

**The PDF file includes:**

Materials and Methods

References

Figs. S1 to S9

**Other Supplementary Material for this manuscript includes the following:**

Table S1. Differentially expressed genes list for WT vs $Tet2^{-/-}$ and $Tet2^{-/-}$ vs $Tet2^{-/-}$; $Id1^{-/-}$ LK cells.

MDAR Reproducibility Checklist

### Materials and Methods

**Histochemical Analysis of Tissue sections.** Sick mice were euthanized with $CO_2$ and organs were removed according to the approved protocol and fixed in 10% neutral buffered formalin for a minimum of five days. After fixation, protocol bones were removed and placed in formic acid decal for 2 days, prior to trimming with the soft tissues. Tissues were processed on an automatic processor and embedding into paraffin blocks. All blocks were sectioned on a manual microtome and placed on charged slides. Slides were dried in an 80°C oven for one hour prior to H&E staining. Hematoxylin and eosin (H&E) staining was performed using the Sakura® Tissue-TekR Prisma™ automated Stainer. The slides were hydrated and stained with commercial hematoxylin, clarifier, bluing reagent and eosin-Y. A regressive staining method was used. This method intentionally overstains tissues and then uses a differentiation step (clarifier/bluing reagents) to remove excess stain. The slides were cover slipped using the Sakura® Tissue-Tek™Glass® automatic cover slipper and dried prior to review. H&E slides were evaluated in a blinded fashion by a pathologist (BK).

**AGX51.** AGX51 was synthesized by Chemical Support Group in the Chemical Biology Laboratory, CCR, NCI, according to the procedure reported by Benezra et al.[1] The compound was characterized by $^{1}H$ NMR, $^{13}C$ NMR, and high-resolution mass spectrometry, all of which agreed with that reported.

**LC/MS-MS (Mass Spectrometry).** BMCs were harvested 12 weeks post competitive transplantation of *WT, Tet2$^{-/-}$, Id1$^{-/-}$* and *Tet2$^{-/-}$; Id1$^{-/-}$* mice. CD45.2$^{+}$/Lin$^{-}$/Sca1$^{+}$/ckit$^{+}$ cells were sorted by FACS and subjected to genomic DNA isolation (Qiagen, AllPrep DNA/RNA Micro

Kit). One micrograms of genomic DNA was digested to nucleoside level using a nucleoside digestion mix (M0649S, New England Biolabs). Mass spectrometry–based quantitation of cytosine, 5mC, and 5hmC was performed as described[2,3]. Briefly, cytosine, 5mC, and 5hmC were quantitated using a Thermo Fisher Scientific Vanquish ultra high-performance liquid chromatography (UHPLC) coupled to a Thermo Fisher Scientific TSQ-Altis tandem mass spectrometer through an electrospray ion source operating in positive ion mode at 3.5 kV. Stock standard solutions were prepared in deionized water at a concentration of 1 mM each. Calibration standard mixtures were prepared at concentrations between 1.0 and 250 μM for dC (2′-deoxycytidine), 0.04 and 10 μM for mdC (5′-methyl-2′-deoxycitine), and 0.002 and 0.5 μM for hmdC (5′hyoxymethyl-2′-deoxycytidine). Linear calibration plots were prepared using concentration versus peak area integration (zero intercept) with a R2 greater than 0.999. By comparing the internal standard normalized peak areas in the digest sample to the corresponding retention times from the calibration standards, the micromolar concentrations of the nucleosides were determined against the standard curve. The molar ratio as a percentage was then calculated as follows: Mol % hmdC = ( [ hmdC ] / ([dC ] + [mdC ] + [hmdC ] ) × 100.

**Lentiviral mediated shRNA knockdown.** Mouse p16 (CDKN2A) (SHCLNG-NM_009877) lentiviral plasmid containing shRNA target sequences were purchased from Sigma-Aldrich. Infectious Lentivirus was generated by transfecting the shRNA constructs and packaging plasmids (PMD2G and pCMV8.74) into 293T/17 cells using LipoD293 (SignaGen, Rockville, MD, USA). Virus containing supernatants were collected 48 hours post transfection. Lentivirus was quantified using Lenti-X Gostix Plus (Takara, #631280), as per the protocol. For knockdown, lineage depleted cells from mouse bone marrow were transduced with shRNA lentivirus by spinoculation. Lineage depleted cells (Depleted for Mac-1, Gr-1, B220, Ter119, CD4, CD8 and Il7R) were isolated from *Tet2*$^{-/-}$*Id1*$^{-/-}$ mice using immune-magnetic bead separation and cultured in Stemspan medium containing mSCF (100 ng/mL), hTPO (100 ng/mL) at $5x10^5$ cells/0.5mL for 12 hours. Twelve hours after culture, the cells were subjected to first round of shRNA mediated lentiviral transduction lentivirus, where lineage depleted cells were spun at 2000 x g for 90 minutes at 37ºC. The cells were then washed and reseeded with fresh complete medium. After 24 hours, a 2nd round of transduction was performed as described above, and the cells were cultured in puromycin (2ug/mL). After 48-72 hours of 2nd lentiviral mediated transduction, the cells were harvested for qRT-PCR and staining with AnnV/7AAD and $C_{12}$FDG for flow cytometry.

**Adenoviral mediated shRNA knockdown.** Mouse Id1(Ad-GFP-U6-m-ID1-shRNA (shADV-261835)) and Scr (Ad-GFP-U6-shRNA, #1122) shRNA silencing adenovirus were purchased from Vector Biosystems Inc. For shRNA mediated Id1 knockdown, lineage depleted cells were isolated from *Tet2*$^{-/-}$ mice bone marrow cells and $5x10^5$ cells/0.5ml were cultured in Stemspan media along with mSCF (100ng/ml) and mTPO (100ng/ml). These cells were transduced with $5ul/5x10^5$ of shId1-GFP or shScr-GFP adenovirus ($10^8$ PFU) on the first and second day of culture. The cells were harvested 5 days post second transduction, stained for p16, AnnV/7AAD and DDAOG and analyzed on flow cytometer.

**Gene Expression Analysis.** RNA was extracted from Donor (CD45.2$^+$)/LSK/ FLT3$^-$/CD150$^+$/CD48$^+$ sorted cells or shRNA transduced cultured cells using the RNeasy Mini Kit (Qiagen, 74104). cDNA synthesis was performed using 1 ug RNA with the iscript cDNA synthesis kit (Bio-Rad, 1708890). Quantitative RT-PCR was performed using the Step-one Plus system with SyBr green (Roche, 04913850001). Amplicons were sequence verified. Genes were normalized to β-actin. Oligonucleotide sequences for the qRT-PCR analysis are as follows:

| **Gene** | **Primer Sequence (5'-3')** |
|---|---|
| p16- Forward Primer | ACT CTT TCG GTC GTA CCC CGA T |
| p16- Reverse Primer | GCA GTT CGA ATC TGC ACC GTA |
| β-actin- Forward Primer | CGG TTC CGA TGC CCT GAG GCT CTT |
| β-actin- Reverse Primer | CGT CAC ACT TCA TGA TGG AAT TGA |

Three biological replicates were performed in each assay with two technical replicates for each sample. Data was analyzed using the standard ∆∆CT method to determine expression changes.

**Karyotyping.** LSM separated BMCs were isolated from *WT, Tet2*$^{-/-}$*, Id1*$^{-/-}$ and *Tet2*$^{-/-}$*; Id1*$^{-/-}$ mice and cultured for 48hrs in IMDM+10%FBS along with mSCF (100 ng/mL), mIL3 (500 ng/mL) and mIL6 (30 ng/mL). Dividing cells were then arrested at metaphase by incubation with Colcemid (KaryoMax Colcemid Solution, Invitrogen, Carlsbad, Calif., USA) (10mg/ml) 3 hrs. before harvest. Cells were collected and treated with hypotonic solution (KCl 0.075 M) for 15 min at 37°C and fixed with methanol: acetic acid 3:1. Slides were prepared, and chromosomal aberrations were analyzed in 15-20 cells per experimental group.

**Bulk RNA-Seq Analysis.** BMCs were harvested 12 weeks post competitive transplantation of *WT, Tet2*$^{-/-}$*, Id1*$^{-/-}$ and *Tet2*$^{-/-}$*; Id1*$^{-/-}$ mice. CD45.2$^{+}$/Lin$^{-}$/Sca1$^{+}$/ckit$^{+}$ cells were sorted by flow cytometry and high-quality RNA was purified using the RNeasy™ Micro Kit (Qiagen, 74004).

The sample was quantified and sequenced on the Illumina NovaSeq 6000 SP sequencer. The HiSeq Real Time Analysis software (RTA 1.18) was used for processing image files, the Illumina CASAVA_v1.8.4 was used for demultiplex and converting binary base calls and qualities to fastq format. The sequencing reads were trimmed for adapters and low-quality bases using Trimmomatic (version 0.30). The trimmed reads were aligned to mouse mm9 reference genome (NCBIM37 /UCSC mm9) and Ensembl annotation version 67 using TopHat_v2.0.8 software. Quantification was carried out with RSEM using the transcriptome bam file created by STAR. Differentially expressed genes (DEGs) were analyzed with Ingenuity Pathway Analysis (IPA) software (Ingenuity Systems, Inc., Redwood City, CA, USA) and Gene Set Enrichment Analysis (GSEA, Broad Institute, Cambridge, MA, USA). For IPA, pathways above a log P-value of 1.33 and a Z-score above 1 were considered statistically significant. For GSEA pathways, Normalized enrichment scores $\geq 1.4$ and an FDR q-value below 0.05 were considered significant. The RNA-Seq data are available in the NCBI's Gene Expression Omnibus (GEO) database (GEO ID: GSE266580).

**Quantification and Statistical Analysis.** Statistical significance was determined using unpaired Student t-tests using Welch's correction when applicable. Kaplan-Meier survival studies were analyzed using the log-rank test. In vivo studies were performed using N=5 mice for each group and were repeated two or more times using an additional N=3-5 mice. *In vitro* studies performed used a minimum N=3 and were repeated 2 or more times. $P \leq 0.05$ was considered statically significant. Error bars portray the standard error of the mean data. *$P \leq 0.05$, **$P \leq 0.01$, ***$P \leq 0.001$.

**Study approval.** Experiments involving the use of mice were approved by the NCI at Frederick Animal Care and Use Committee in accordance with the eighth edition "Guide for the Care and Use of Laboratory Animals."

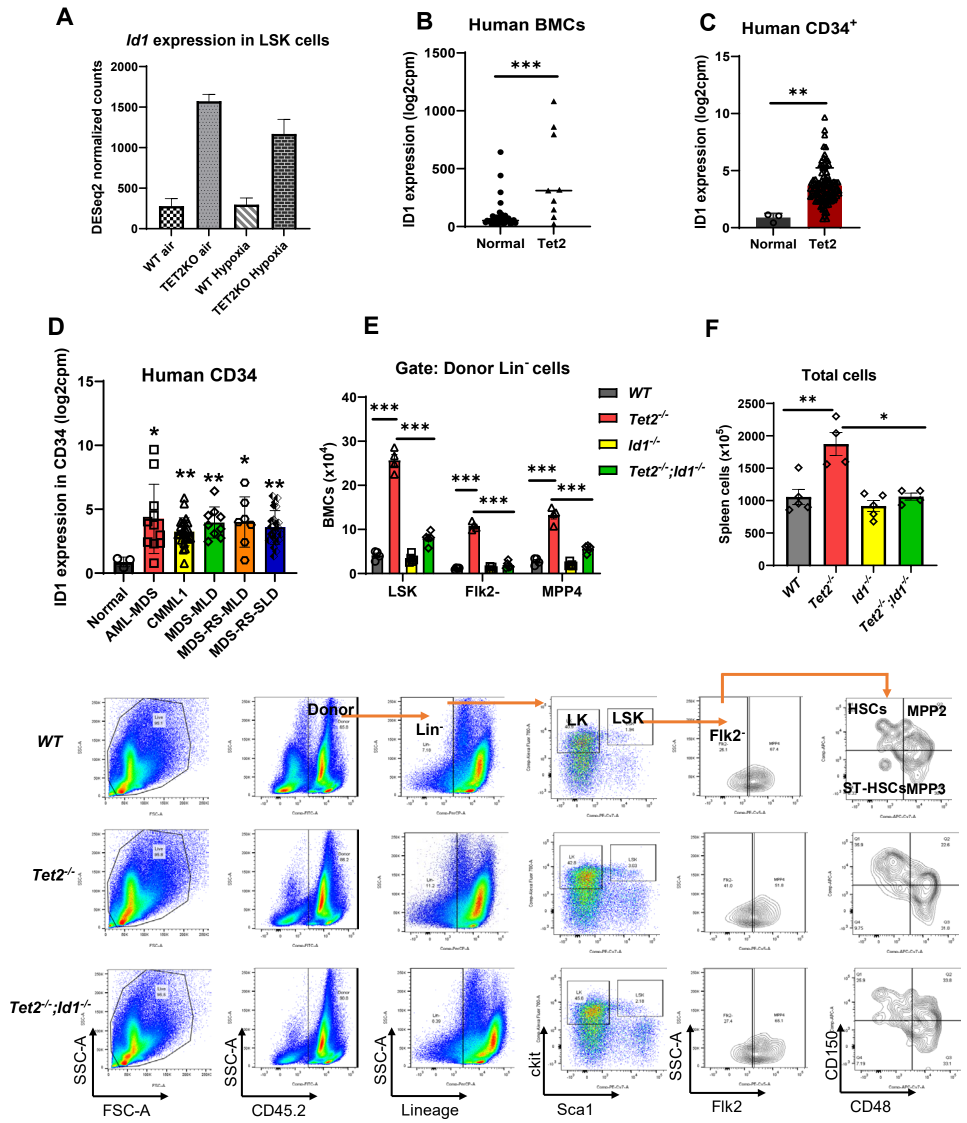


**Fig. S1. *Id1* expression is upregulated in *Tet2^-/-^* HSPCs.** (A) Expression of *Id1* in *WT* and *Tet2*$^{-/-}$ LSK cells from published RNA-seq data set[1]. (B) *ID1* expression in the human unfractionated, (C) CD34$^+$ BMCs from AML patient samples with *Tet2* mutations and (D) CD34$^+$ BMCs from patient with MDS and CMML. (E) Total number of donor derived LSK, LSK/Flk2$^-$ and MPP4 cells in competitive BMT recipient mice. (F) Total number of donor derived cells in spleens of competitive BMT recipient mice. (G) Flow cytometry gating strategy for the analysis of donor derived HSPCs in competitive BMT recipients. Data are presented as the mean ± SEM, and comparisons between mean values of groups were evaluated using an unpaired, 1-tailed Student's t test. *$P \leq 0.05$, **$P \leq 0.01$ and *** $P \leq 0.001$. AML-MDS: AML with myelodysplasia-related changes, CMML1: chronic myelomonocytic leukemia 1, MDS-MLD: MDS with multilineage dysplasia, MDS-RS-MLD: MDS with ring sideroblasts with multilineage dysplasia, MDS-RS-SLD: MDS with ring sideroblasts with single lineage dysplasia.

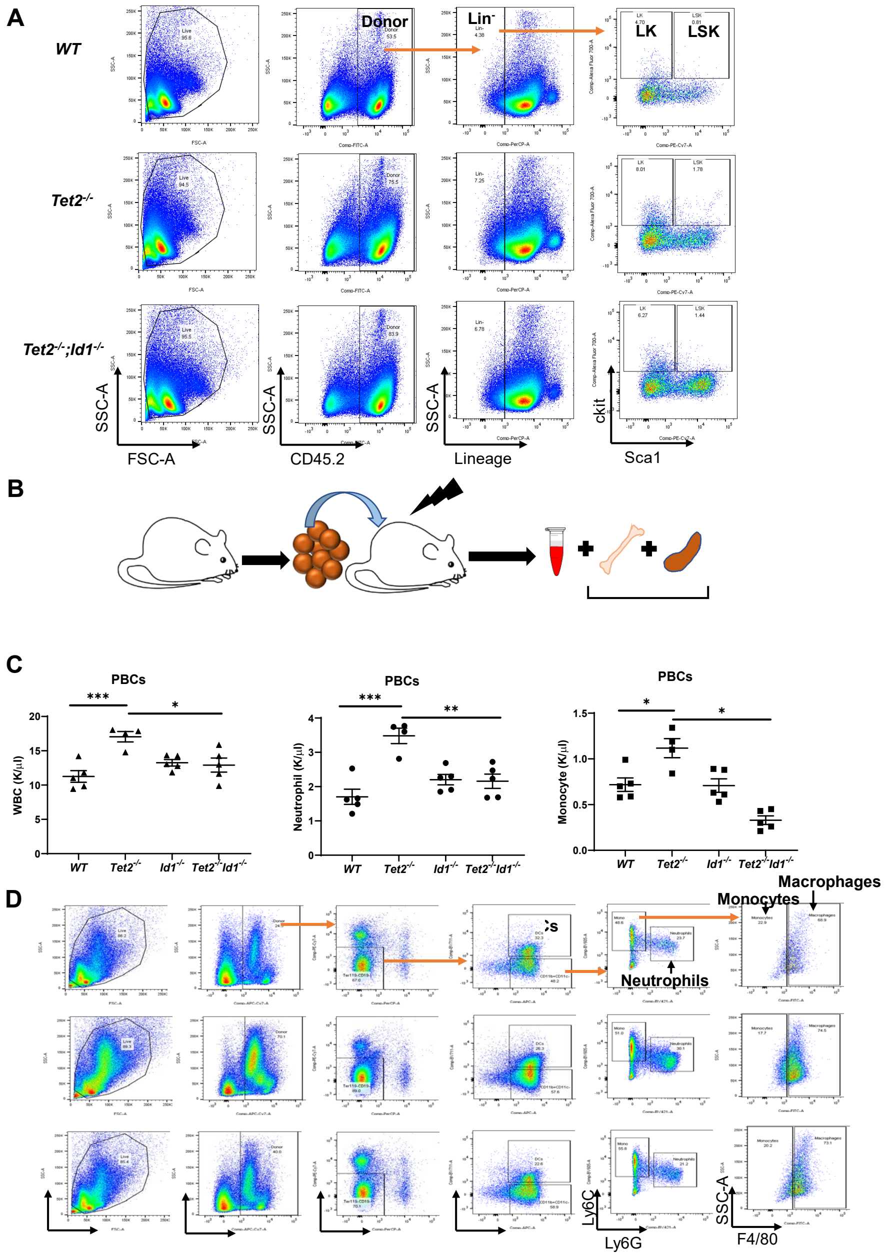


**Fig. S2. Loss of *Id1* expression in *Tet2*$^{-/-}$ HSPCs reduces and rescues myeloid skewing.** (A) Flow cytometry gating strategy for analysis of spleen HSPCs in competitive BMT recipient mice. (B) Summary of experimental design for analysis of myeloid skewing in primary BMT recipient mice. (C) CBC analysis of PBCs from noncompetitive BMT recipient mice after 12 wks. (D) Flow cytometry gating strategy for analysis of donor derived neutrophils, dendritic cells (DCs), monocytes, macrophages in competitive BMT recipient mice. Data are presented as the mean ± SEM, and comparisons between mean values of groups were evaluated using an unpaired, 1-tailed Student's t test. $^*P \leq 0.05$, $^{**}P \leq 0.01$ and $^{***} P \leq 0.001$

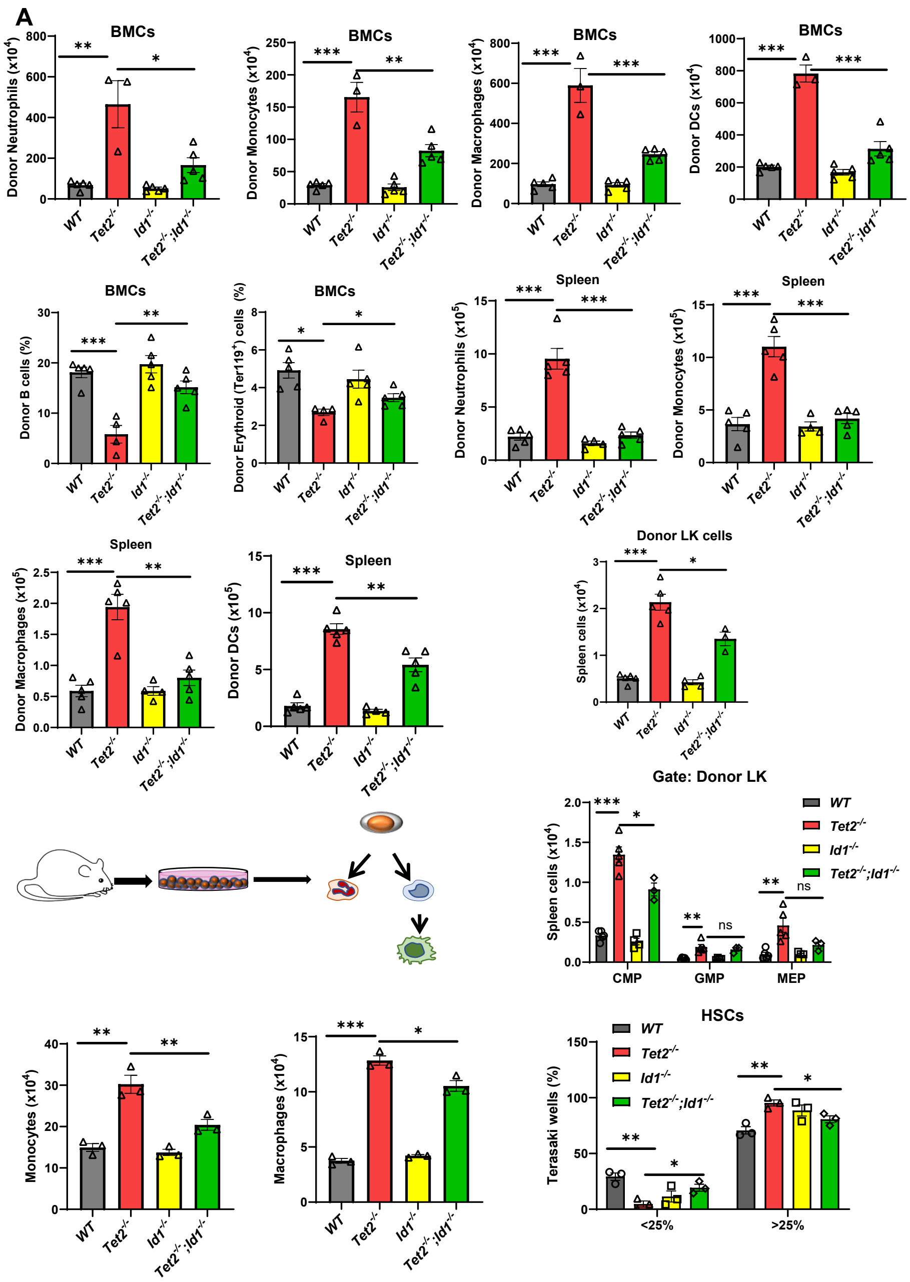


**Fig. S3. Ablation of *Id1* in *Tet2*$^{-/-}$ HSPCs reduces myeloid skewing in the BM and spleen of BMT recipient mice.** (A) Total number of donor neutrophils, monocytes, macrophages, and dendritic cells in the BM of competitive BMT recipient mice. (B) Frequency of donor derived $CD19^{+}$ B cells and erythroid progenitors (CD45+/$Ter119^{+}$) in the BM of competitive BMT recipient mice. (C) Total number of donor derived neutrophils and monocytes and (D) macrophages and dendritic cells in spleens of competitive BMT recipient mice. (E) Summary of experimental procedure to evaluate the differentiation potential of Lin- BMCs *in vitro*, and total number of monocytes and macrophages after 6 days of *in vitro* culture. (F) Total number of donor derived LK cells and (G) CMP, GMP and MEP cells in spleen cells from competitive BMT recipient mice. (H) Single cell growth assay of FACS sorted HSCs in Terasaki plates determined by the percentage of wells with <25% or >25% confluency after seven days of culture. Data are presented as the mean ± SEM, and comparisons between mean values of groups were evaluated using an unpaired, 1-tailed Student's t test. *$P \leq 0.05$, **$P \leq 0.01$ and *** $P \leq 0.001$.

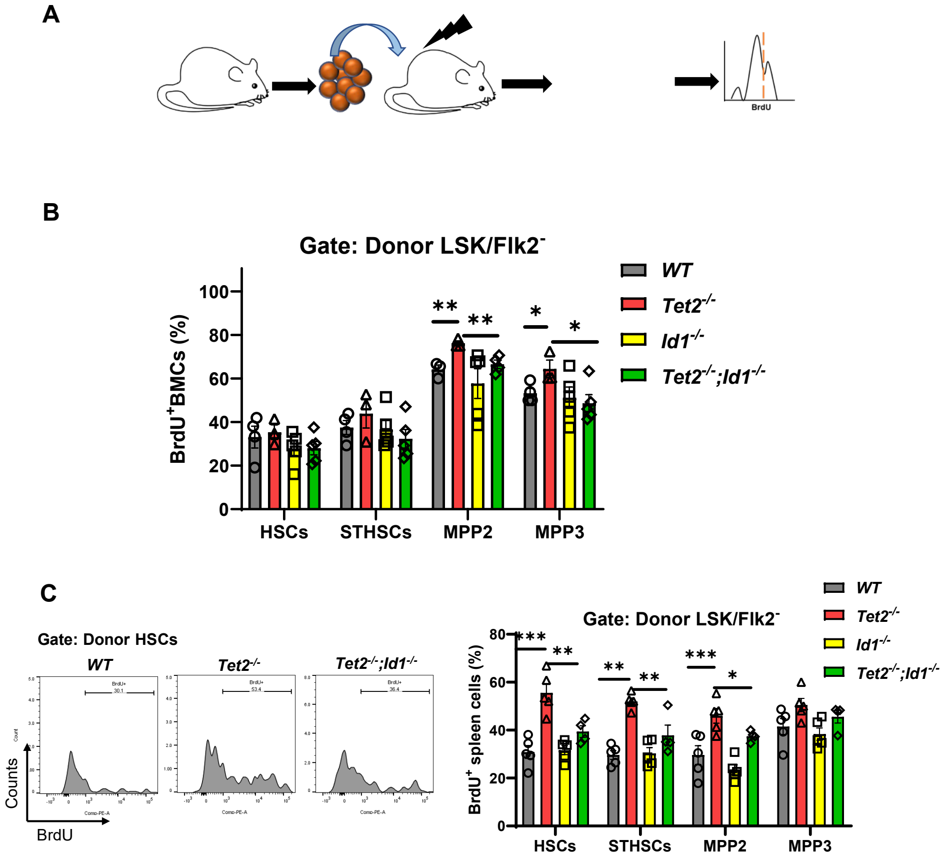


**Fig. S4. Ablation of *Id1* expression in *Tet2*$^{-/-}$ HSPCs reduces cell cycling and proliferation of spleen cells from competitive BMT recipient mice. (A)** Experimental design to evaluate cell cycling and proliferation of HSPCs in competitive BMT recipient mice. (B) Frequency of BrdU$^{+}$ donor BM HSPCs in competitive BMT recipient mice. (C) Flow cytometry histograms of BrdU stained donor spleen HSCs, and the percentage of BrdU$^+$ donor spleen HSPCs in competitive BMT recipient mice. Data are presented as the mean ± SEM, and comparisons between mean values of groups were evaluated using an unpaired, 1-tailed Student's t test. *$P \leq 0.05$, **$P \leq 0.01$ and *** $P \leq 0.001$

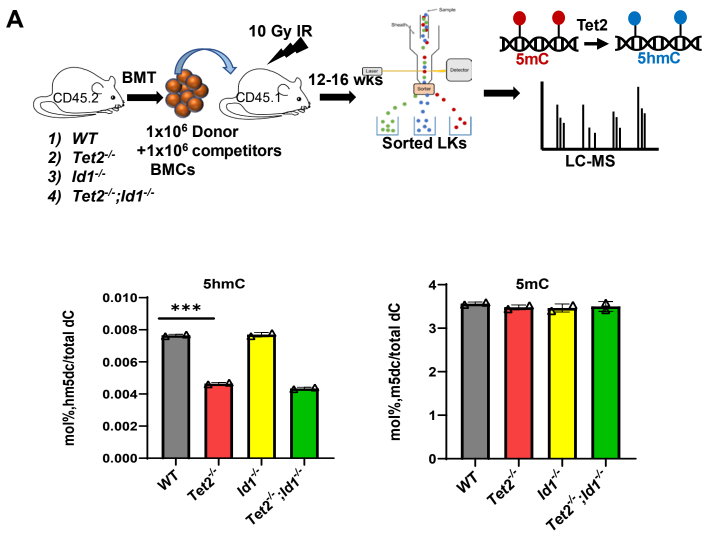

**Fig. S5. *Id1* expression does not affect the levels of 5hmC in *Tet2*$^{-/-}$ HSPCs but influences the *Tet2*$^{-/-}$ transcriptome. (A)** Summary of procedure to measure 5mC and 5hmC levels in donor LK cells via Liquid Chromatography-Mass Spectrometry (LC-MS) from competitive BMT recipient mice. (B) Quantification of 5hmC and 5mC levels in donor LK cells from competitive BMT recipient mice. Data are presented as the mean ± SEM, and comparisons between mean values of groups were evaluated using an unpaired, 1-tailed Student's t test. *** $P \leq 0.001$

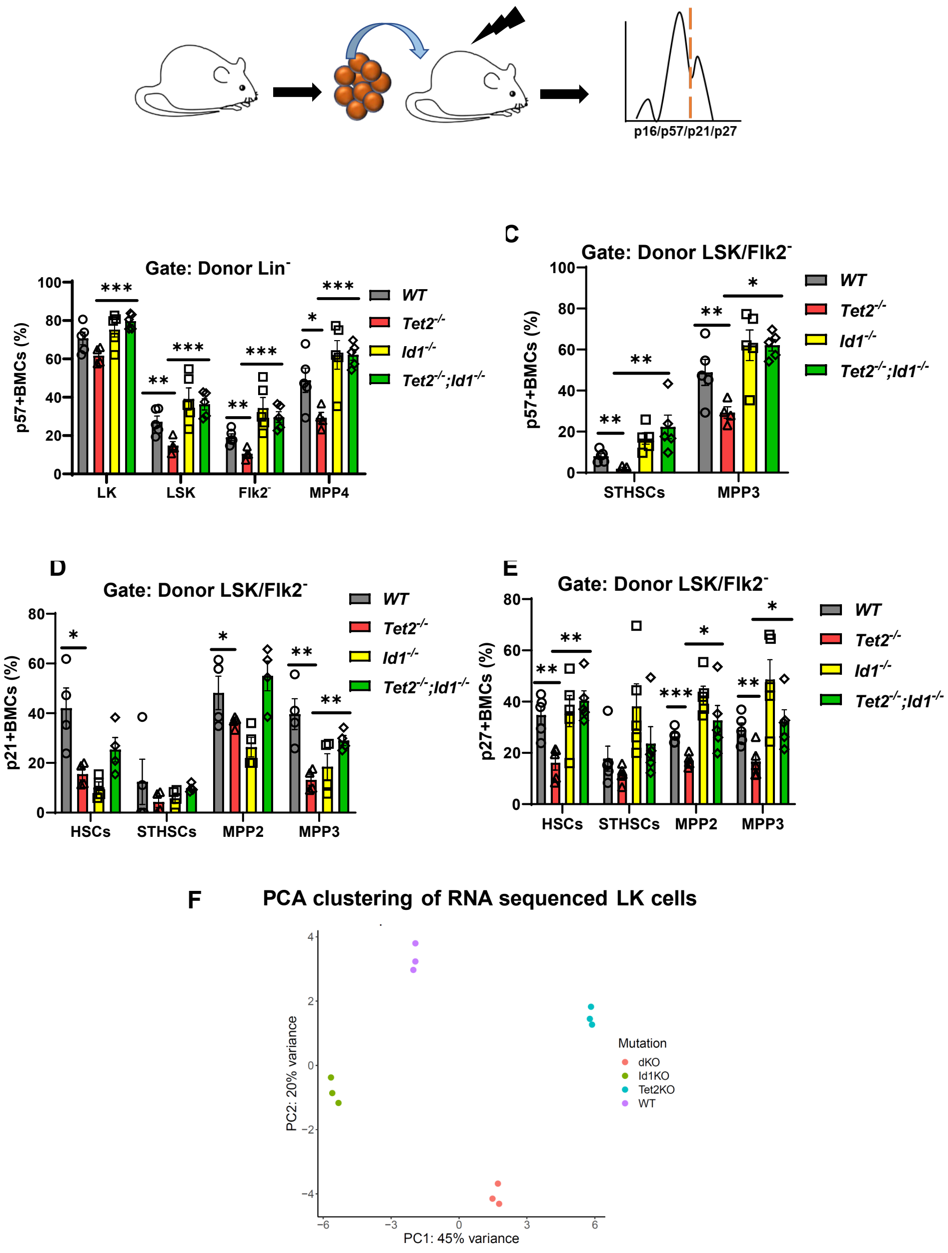


**Fig. S6. Loss of *Id1* increases cyclin-dependent kinases inhibitor gene expression in *Tet2*$^{-/-}$ BMCs**. (A) Summary of experimental design to evaluate cyclin dependent kinase inhibitor (CDKI) expression in donor derived HSPC from competitive BMT recipient mice. Frequency of donor HSPCs that express (B-C) p57, (D) p21 and (E) p27 in BMCs from competitive BMT recipient mice by flow cytometry. (F) Principal component analysis (PCA) plot of FACS sorted donor LK from competitive BMT recipient mice. Data are presented as the mean ± SEM, and comparisons between mean values of groups were evaluated using an unpaired, 1-tailed Student's t test. *$P \leq 0.05$, **$P \leq 0.01$ and *** $P \leq 0.001$

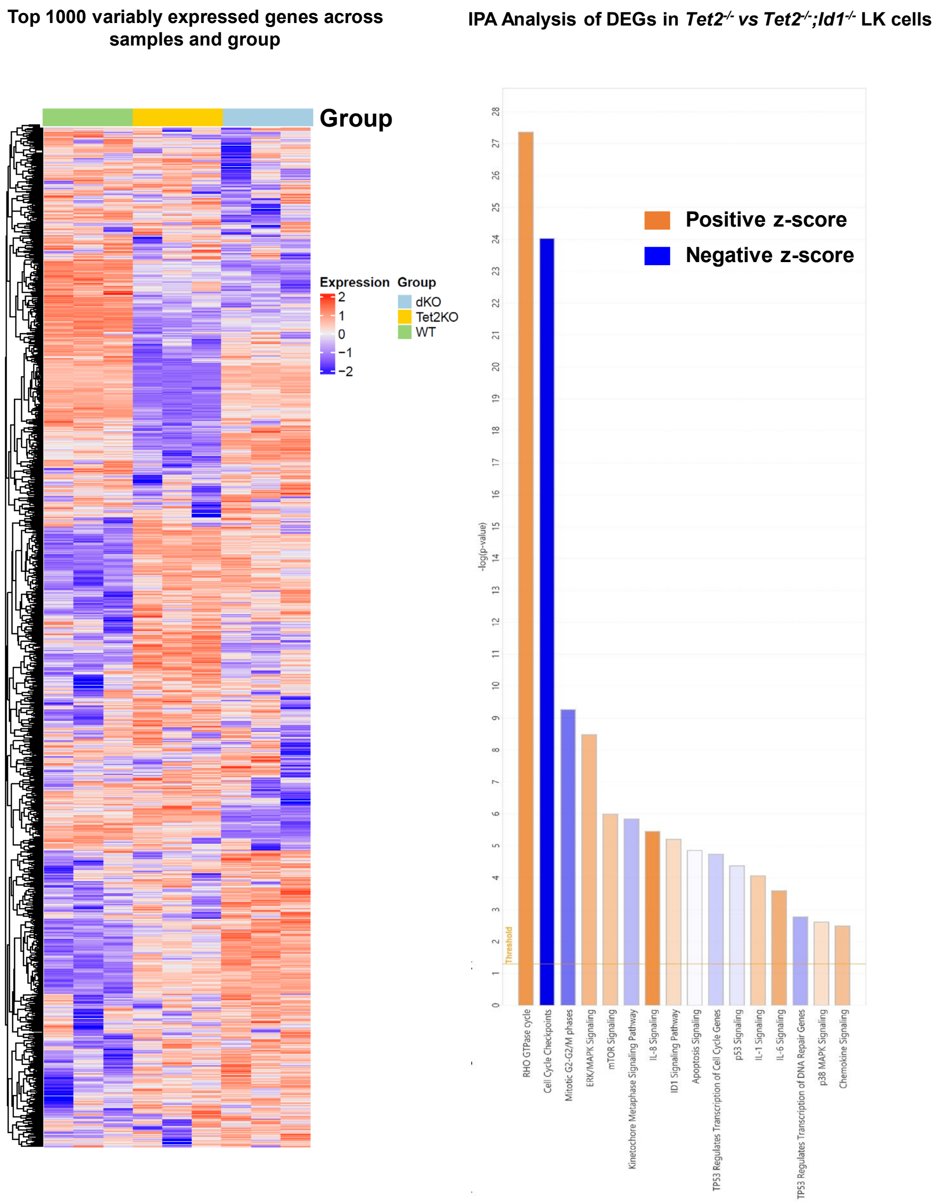


**Fig. S7. RNA-sequence and differential gene expression (DEG) analysis of donor *WT*, *Tet2*$^{-/-}$ and *Tet2*$^{-/-}$; *Id1*$^{-/-}$ LK cells**. (A) Heatmap of top 1000 variably expressed genes in WT and *Tet2*$^{-/-}$, and *Tet2*$^{-/-}$; *Id1*$^{-/-}$ donor LK cells from competitive BMT recipient mice. (B) IPA analysis of differentially expressed genes with a cut-off of adjusted p value (q value) of $\leq 0.05$, in donor *Tet2*$^{-/-}$ versus *Tet2*$^{-/-}$; *Id1*$^{-/-}$ LK cells from primary BMT recipient mice. The significant canonical pathways affected by this analysis are indicated on the y-axis. Bar colors indicate predicted pathway activation (orange), and predicted inhibition (blue) (z-score). The significance values for the canonical pathways are calculated by Fisher's exact test (-log of p-value).

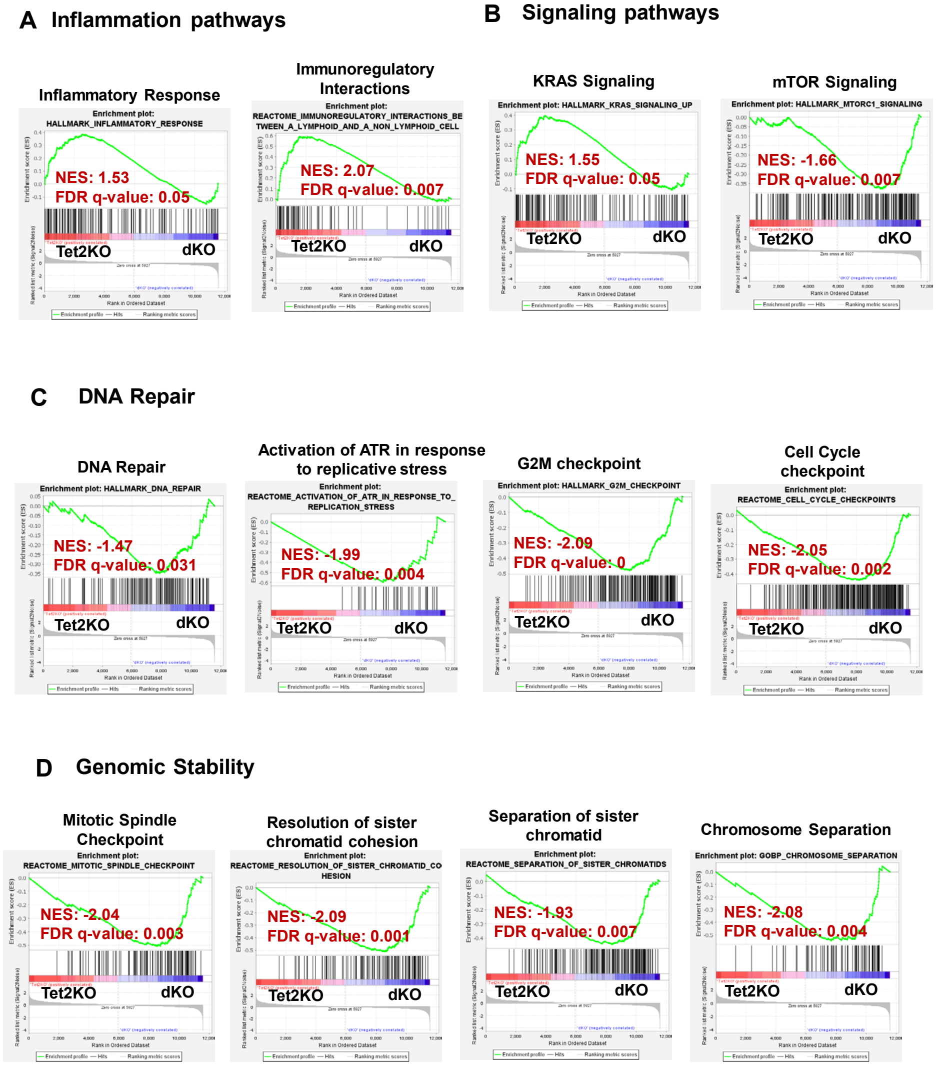


**Fig. S8. GSEA analysis of differentially expressed genes in *Tet2*$^{-/-}$ and *Tet2*$^{-/-}$; *Id1*$^{-/-}$ LK cells.**

GSEA analysis of differentially expressed genes in donor *Tet2*$^{-/-}$ versus *Tet2*$^{-/-}$; *Id1*$^{-/-}$ LK BMCS from competitive BMT recipient mice shows (A) reduced expression of inflammatory response and immunoregulatory interactions genes in *Tet2*$^{-/-}$; *Id1*$^{-/-}$ LK cells, (B) reduced expression of genes regulating cell signaling pathways in *Tet2*$^{-/-}$; *Id1*$^{-/-}$ LK cells, (C) increased expression of genes involved in the DNA repair pathway, (D) increased expression of genes involved in genomic stability in *Tet2*$^{-/-}$*; Id1*$^{-/-}$ donor LK cells. This analysis was performed using molecular signature database: Hallmark, Reactome and Gene Ontology Biological Processes (GOBP) gene sets.

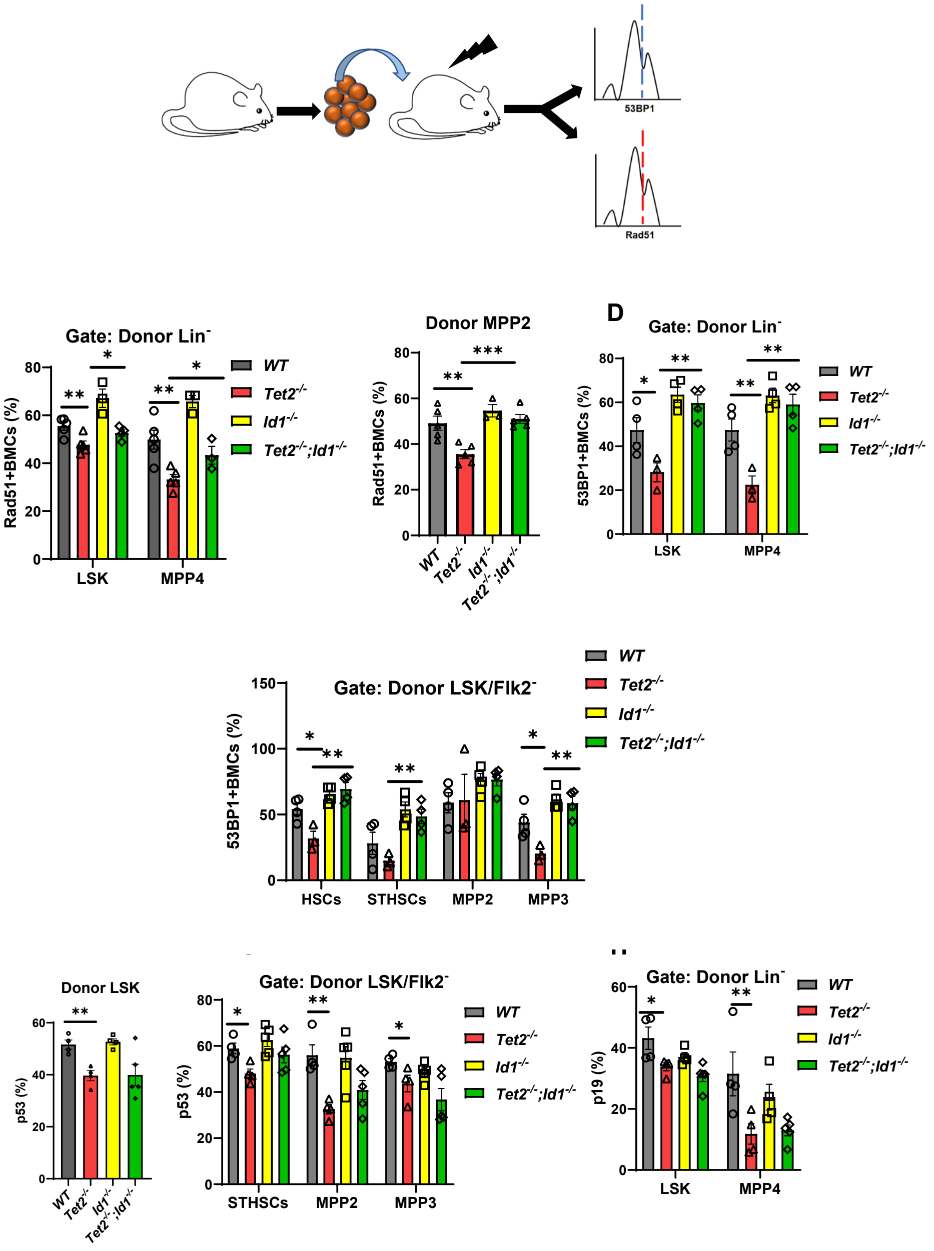


**Fig. S9. *Id1* ablation leads to increased DNA repair protein expression in *Tet2*$^{-/-}$ BMCs**. (A) Summary of procedure to determine the expression of DNA damage response genes, 53BP1 and Rad51, in HSPCs from competitive BMT recipient mice. (B) Frequency of Rad51 expressing donor LSK and MPP4 and (C) MPP2 BMCs in competitive BMT recipient mice. (D) Percentage of 53BP1 expressing donor LSK, MPP4 and (E) HSPCs in BMCs of competitive BMT recipient mice. (F) Percentage of p53 expressing donor LSK and (G) HSPCs in BMCs of competitive BMT recipient mice. (H) Frequency of p19 expressing donor LSK and MPP4 BMCs in competitive BMT recipient mice. Data are presented as the mean ± SEM, and comparisons between mean values of groups were evaluated using an unpaired, 1-tailed Student's t test. $*P \leq 0.05$, $**P \leq 0.01$ and $***P \leq 0.001$
